# Parallel evolution of mutational fitness effects over 50,000 generations

**DOI:** 10.1101/2022.05.17.492023

**Authors:** Anurag Limdi, Siân V. Owen, Cristina M. Herren, Richard E. Lenski, Michael Baym

## Abstract

As evolving populations accumulate mutations, the benefits and costs of subsequent mutations change. As fitness increases, the relative benefit of new mutations typically decreases. However, the question remains whether deleterious mutations tend to have larger or smaller costs as a population adapts; theory and experiments provide support for both conflicting hypotheses. To address this question, we compared the effects of insertion mutations in every gene in *Escherichia coli* between ancestral and 12 independently derived strains after 50,000 generations in a uniform environment. We found both increases and decreases in the fitness costs of mutations, leaving the overall distribution of effects largely unchanged. However, at the extreme, more genes became essential over evolution than vice versa. Both changes in fitness effects and essentiality evolved in parallel across the independent populations, and most changes were not explained by structural variation or altered gene expression. Thus, the macroscopic features of the local fitness landscape remained largely unchanged, even as access to particular evolutionary trajectories changed consistently during adaptation to the experimental environment.

**One Sentence Summary:** Limdi et al. report parallel changes in the cost of mutations in replicate lineages of a decades-long *E. coli* evolution experiment.

## Main Text

Evolution is a local process. As an asexual population evolves, natural selection can only act upon new genotypes that arise in the mutational neighborhood of the resident population. With every accumulating mutation, the accessibility and effects of subsequent mutations can change through genetic interactions (*1–3*). Therefore, the local mutational neighborhood may look increasingly different from the neighborhood of the ancestral genotype over long periods of evolution. *Adaptability*, namely the availability and benefits of new mutations in the local neighborhood, tends to decrease over time in microbial populations as they evolve in a constant environment (*4–8*). What happens to *mutational robustness*, namely the prevalence and steepness of mutational paths leading to lower fitness, is less clear.

Previous experiments have yielded conflicting results about how robustness changes over evolution. In viruses, robustness to mutations and environmental perturbations has been reported both to increase and decrease during evolution (*9, 10*). In evolving yeast populations, robustness to insertion mutations has either declined or remained unchanged depending on the environment (*11*).

Theory does not resolve this conflict. Selection can favor mechanisms conferring increased robustness to mutations, especially at high mutation rates and in large populations (*12–15*). However, recent theoretical work suggests that widespread interactions between mutations accumulated over evolution may lead to an emergent pattern of increasing cost epistasis (*16*): on average, mutations are more detrimental on fitter genetic backgrounds, an idea supported by experiments with diverse yeast strains (*17*). However, theory is limited to broad statistical properties of the mutational neighborhood, which can obscure changes in the importance of specific genes.

The most extreme changes are found in essential genes, whose loss renders an organism inviable. Prior work has shown that essentiality is not a static property of genomes (*18*); the set of essential genes can vary greatly between species and even strains of the same species (*19, 20*). For instance, about a third of the essential genes in *E. coli* are nonessential in *Bacillus subtilis*, and vice versa (*21*). Gene essentiality is also malleable over shorter timescales; in *Saccharomyces cerevisiae* and *Staphylococcus aureus*, many essential genes become nonessential following selection for suppressors (*22, 23*), and horizontal gene transfer via prophages impacts essentiality of core genes in *E. coli* (*20*). To what extent gene essentiality changes in the absence of these driving forces remains unclear.

To investigate changes in robustness, we turned to the Long-Term Evolution Experiment (LTEE), in which twelve populations of *E. coli* have been serially propagated in a glucose-limited minimal medium (*24*) for over 75,000 generations. If there is a systematic evolutionary trend in robustness, then the large number of generations as well as the lack of recombination, horizontal gene transfer, and environmental change make this system ideally suited to detect it. In both the preserved ancestral strains and after 50,000 generations we measured the effects on fitness of disrupting every gene with an inserted transposon. Such insertions typically lead to losses of function – by their nature, spontaneous loss-of-function mutations occur readily, and so our approach effectively surveys a large part of the fitness landscape accessible by single-step mutations. We measured the fitness of the mutants by sequencing, preserving the identity of each mutant and allowing interrogation not just of overall trends, but of the specific changes in the locally accessible landscape.

## Results

### Simultaneous parallel fitness measurements of thousands of insertion mutants

To measure how robustness changed during evolution, we used a suicide-plasmid delivery system to construct high-coverage transposon libraries in the LTEE ancestors (REL606 and REL607) and a clone isolated from each population (Ara+1 to Ara+6 and Ara−1 to Ara−6) at 50,000 generations. The *mariner*-derived transposon, ∼2.2kb in length, inserts into TA dinucleotides, providing over 200,000 possible insertion sites in the ancestral chromosome. We propagated the mutant libraries at 37C in Davis minimal media with 25 mg/L glucose (DM25) for four days, diluting the cultures 1:100 in fresh medium each day. These conditions are the same as those used in the LTEE, and the bacteria have log_2_(100) ≈ 6.64 generations per day. The cost or benefit of each mutation determines the expected rate at which it grows relative to the rest of the population. The change in the abundance of each mutation can be used to infer its relative fitness effect using insertions in non-functional pseudogenes, in the same transposon library, as putative neutral markers (outlined in Methods). Hence, relative fitness effects of mutations can be compared across the LTEE strains.

Measuring fitness effects requires precise quantification of the changes in each mutant’s abundance. To that end, we attached unique molecular identifiers (UMIs) to individual molecules in transposon insertion libraries, and quantified abundance by sequencing across transposon-genome junctions at >100X depth. We observed transposon insertions throughout the genome, with >100,000 unique insertions in all libraries, disrupting over 83% of the genes, with >95% overlap in genes disrupted in the ancestral and evolved libraries (Fig. S1).

We then estimated the fitness effect of disrupting each gene by averaging the fitness effects over all the TA insertion sites in the interior region of the gene (Fig. 1). Transposon insertions do not always lead to loss of function, so we excluded genes that appeared non-essential in the *E. coli* K-12 TraDIS dataset (generated using the Tn5 transposon) due to insertions at C-termini (e.g., *spoT*) or insertions in non-essential gene domains (e.g., *ftsK, ftsN*) (*25*). The resulting fitness estimates were highly reproducible between technical replicates, and consistent with independent estimates obtained by pairwise competitions between engineered deletion mutants and their unmutated parents (Fig. S2, Table S1).

**Fig. 1:**
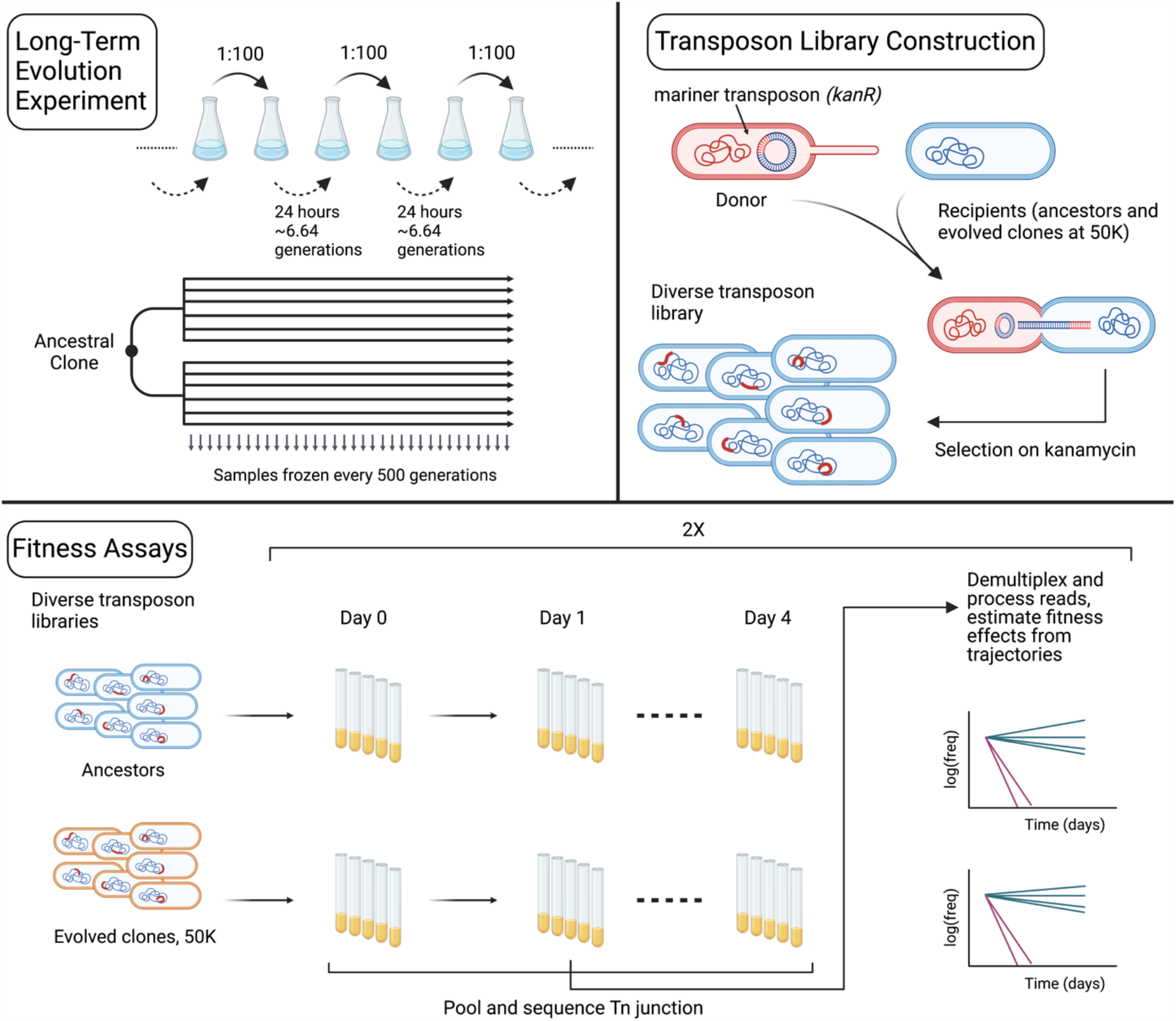
Schematic representation of mutagenesis and fitness assay pipelines. The Long-Term Evolution Experiment (LTEE) is an ongoing experiment in which 12 populations of *E. coli* evolve in and adapt to a glucose-limited minimal medium. Here, we created transposon libraries in the LTEE ancestors and clones sampled from each population at 50,000 generations. We conjugated the strains with *E. coli* MFDλpir-pSC189, which carries a mariner transposon with a *kanR* resistance gene, and we selected transconjugants on medium containing kanamycin. We then propagated the resulting insertion libraries for four days in the same minimal medium as used in the LTEE, and we quantified the abundance of mutants over time using sequence data. We measured the fitness effects of insertions in each gene in the genome based on changes in the relative abundance of the mutants. These measurements were highly reproducible and consistent with fitness effects based on pairwise competitions (Fig. S2). The abundance trajectories shown here are illustrative examples. Figure created with biorender.com.

### No systematic changes in the distribution of fitness effects

We compared the overall distribution of fitness effects (DFEs) in the mutant populations derived from the two LTEE ancestors and 12 evolved clones. We excluded two populations derived from evolved clones from further analyses because their fitness measurements were unreliable for technical reasons, and therefore not comparable to the ancestor. In Ara+4, the within-gene measurement variability for fitness was extremely high, and the correlation between technical replicates was poor (Fig. S3A). In Ara–2, a few insertion mutations increased rapidly, outcompeting other mutations (Fig. S3, B and C), which made the measurements unreliable and systematically biased (see Supplementary Text 1 for more details).

We found that most mutations are nearly neutral (within ∼2-3% of neutrality, depending on the strain), but in all cases with a somewhat heavier tail of deleterious mutations than beneficial mutations (Fig. 2A), consistent with previous results (*26–28*). The aggregate DFEs for the ancestors and evolved lines were nearly identical, except for an excess of mutations that are beneficial (*s* > 0.03, an effect reliably distinguishable from measurement noise) in the ancestral vs evolved backgrounds (0.9% vs 0.5% of all mutations, respectively; Fig. 2B, note the logarithmic scaling). This difference in the supply of beneficial mutations and its evolutionary significance are examined in depth in the companion study by Couce et al (*29*).

**Fig. 2:**
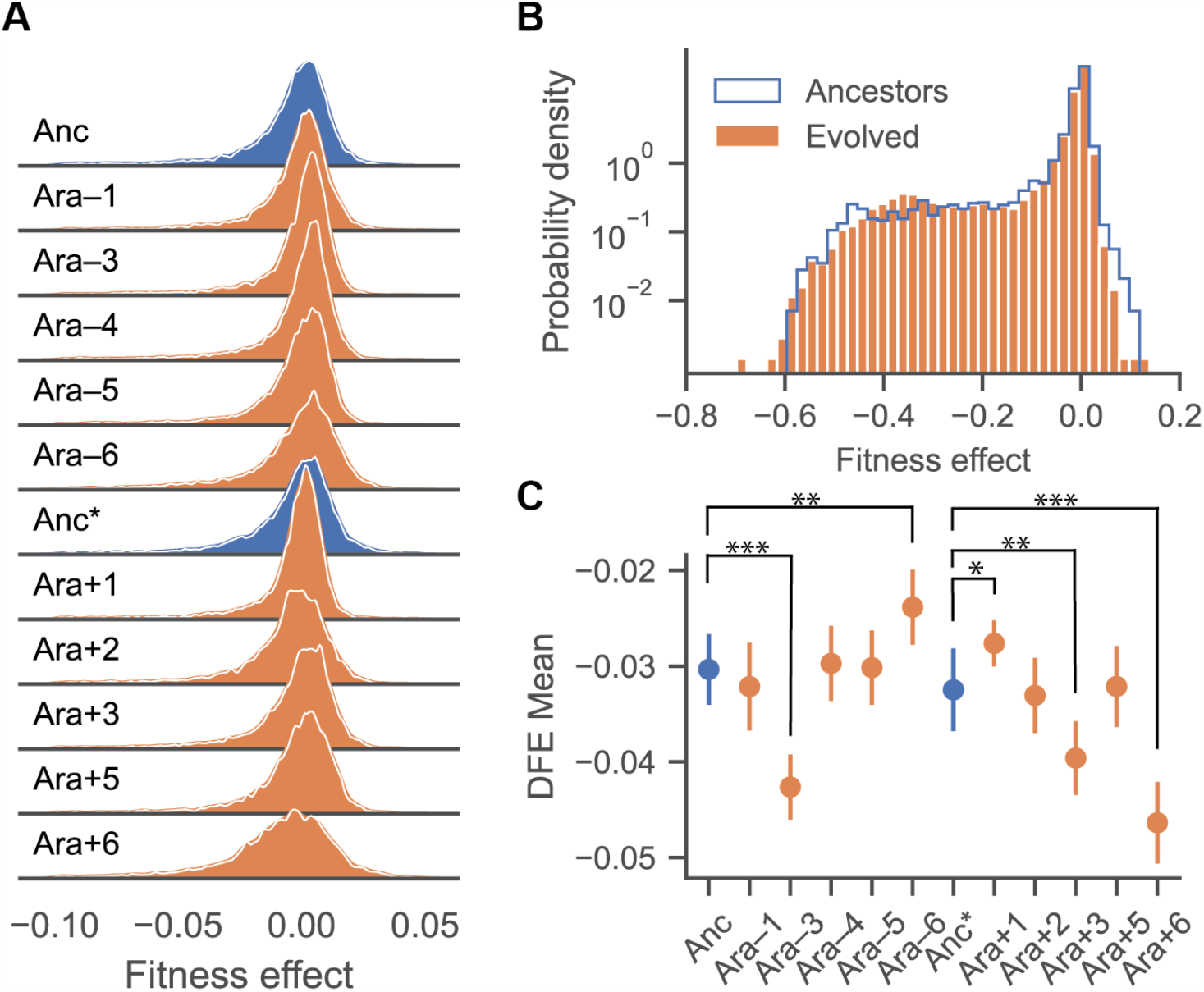
The overall distribution of fitness effects is largely unchanged after 50,000 generations. **(A)** Ridge plot of the overall distribution of fitness effects (DFE) in the LTEE ancestors (Anc: REL606, Anc*: REL607) (blue), which differ by a neutral marker, and 50,000-generation clones sampled from each population (orange). We excluded two strains (Ara–2 and Ara+4) from further analyses (see text; Fig. S3). The histograms were smoothed using kernel density estimation and are shown with a linear y-axis. DFEs are only shown for fitness effects ranging from –0.1 to 0.05, as the density outside these regions is very low. **(B)** Comparison of the aggregated DFEs of the ancestral and evolved strains. Here the histograms are plotted with a logarithmic y-axis to show more clearly the deleterious and beneficial tails of the DFEs. **(C)** Means of the DFEs: error bars indicate the 95% confidence interval in the estimate of means given the associated measurement noise in the bulk fitness assays. Statistically significant differences between the evolved lines and ancestors after Bonferroni correction for multiple tests are indicated (*Z*-test,; ∗∗∗ *p* < 0.001, ∗∗ 0.001 < *p* < 0.005, ∗ 0.005 < *p* < 0.05).

There was no systematic directional trend in how the means of the DFEs changed during evolution (*t*-test based on population means: *p* = 0.37). While the mean fitness effect differed significantly between the ancestor and several evolved lines considered individually (Fig. 2C), these differences vary in their direction (2 evolved clones had higher means than the ancestor, and 3 evolved clones had lower means), and they are primarily driven by noisy measurements at the deleterious tail (Fig. S4). Therefore, robustness measured as the overall mean of the DFE of gene disruption did not systematically change during the 50,000 generations of adaptation.

### Parallel changes in fitness effects over evolution

We then examined how the fitness effects of the same insertion mutations varied between the ancestor and the evolved strains. We restricted analysis to gene disruptions with fitness effects *s* > –0.3 in both the ancestor and evolved strain, as measurements of highly deleterious effects are more prone to measurement noise. The fitness effects of many mutations differed between the ancestral and evolved strains, with some becoming more deleterious and others less deleterious (Fig. 3A). Depending on the evolved strain, between 3% and 6% of the insertion mutations had significantly different fitness effects from those in the ancestor (Fig. 3B), and 13% of the insertion mutations had differential fitness effects in at least one evolved strain.

**Fig. 3:**
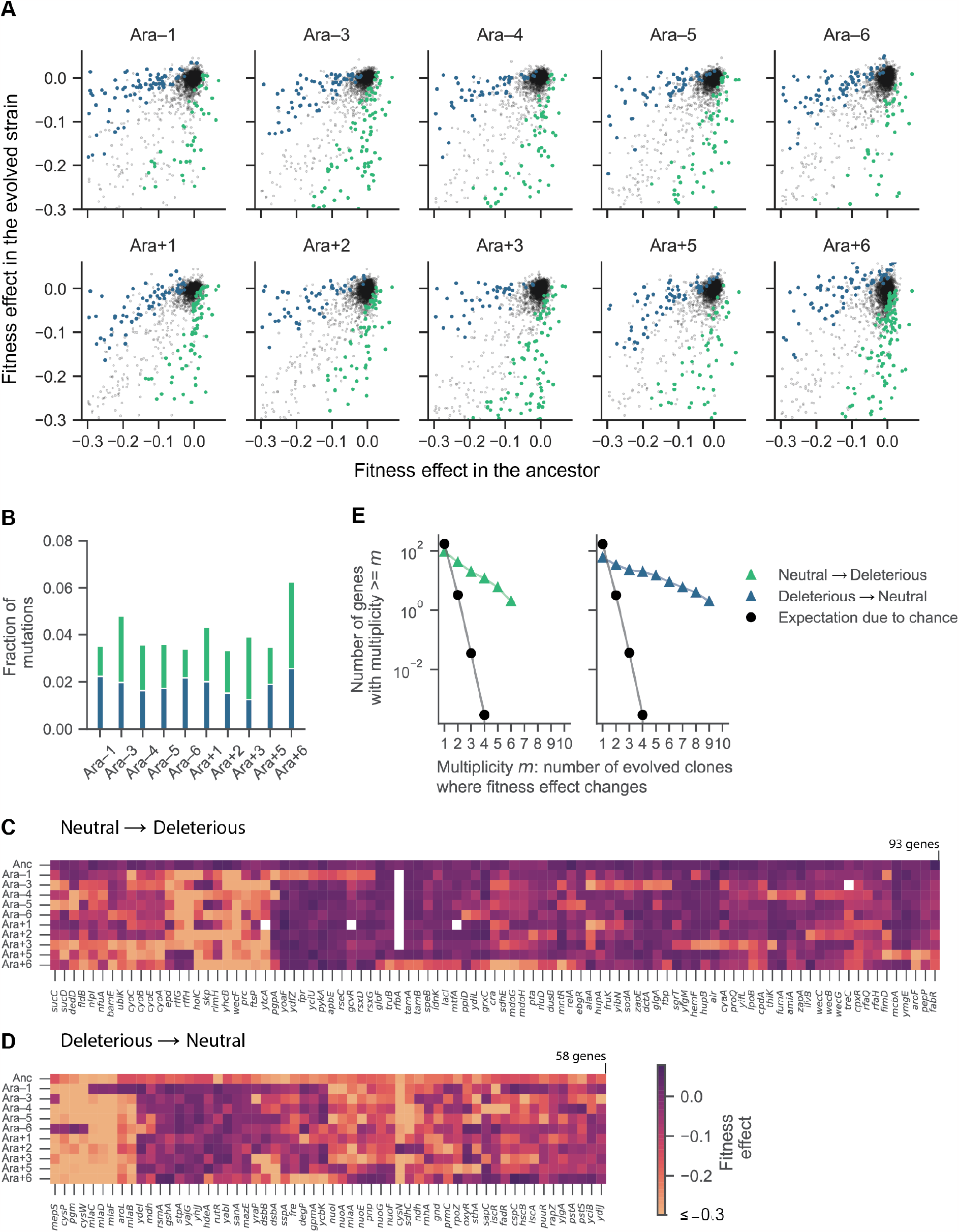
Extensive and parallel changes in fitness effects of gene disruption over evolution. **(A)** Pairwise comparison of fitness effects of mutations in nonessential genes (*s* > –0.3) between the ancestor (REL606) and each evolved strain. Blue: more deleterious in the ancestor, green: more deleterious in the evolved strain, Bonferroni corrected *p* value < 0.05 (two-tailed *Z*-test). **(B)** Fraction of mutations (with *s* >–0.3 in both the ancestor and the evolved strain) with significant differences in fitness effects between the ancestor and each evolved clone (Bonferroni corrected *p*-values < 0.05). **(C, D)** Clustered heatmaps showing fitness effects (scale at right) of gene disruptions that became neutral (*s* > –0.05) or deleterious (–0.3 < *s* < –0.15) in at least one 50,000-generation strain. Genes that were deleted during evolution are shown in white. Genes with mutations conferring fitness effects below –0.3 (the threshold for essentiality) were set to –0.3 for the clustering and visualization. **(E)** Parallel changes in fitness effects. We estimated the expected number of parallel changes from chance alone by shuffling the fitness profiles 10,000 times and counting how often the same genes had parallel changes (neutral to deleterious or deleterious to neutral) in at least *m* populations. The expectation is an average over 10,000 simulations, and therefore it can be < 1.

Comparing genes with fitness effects that changed significantly over evolution, we observed significant parallelism across the independent lineages. We first examined this possibility through hierarchical clustering of mutations that were neutral in the ancestor (*s* > –0.05) and deleterious in an evolved strain (–0.3 < *s* < –0.15), and vice versa (Fig. 3, C and D). While many such changes were specific to individual lineages, many others occurred in parallel across multiple lineages. To assess whether the observed parallelism was greater than expected by chance, we compared the two complementary cumulative distributions of differentially effects of gene disruptions in multiple lineages against a null distribution, which we generated by shuffling the fitness profiles of each population 10,000 times. Both the neutral-to-deleterious and deleterious-to-neutral transitions occurred in parallel much more often than expected by chance (Fig. 3E). This outcome was insensitive to the chosen cutoff values (Fig. S5). These parallel changes across the evolutionary replicates imply that these changes in fitness effects are a result of selection.

### Parallel changes in gene essentiality over evolution

Gene essentiality, in the sense of lethality or an absolute inability to replicate, is often difficult to distinguish from extreme growth defects. For this analysis, we define a gene as differentially essential between the ancestor and an evolved clone if (i) the fitness effect of disruption *s* > –0.15 in one strain and *s* < –0.3 in the other, or (ii) mutants were absent in the library prior to selection in DM25, suggesting that the gene was essential in LB (see Methods for details). This approach ensured that small changes in fitness effects (say from –0.31 to –0.29) were not counted as changes in essentiality. Further, our choice of *s* < –0.3 emerged from simulated fitness assays, which indicated that mutations with deleterious effects of this magnitude or larger could not be reliably distinguished from lethality (Fig. S6). Using the cutoff *s* < –0.3, we detected 557 genes that were essential in DM25 (see Methods).

We found genes that went from nonessential to essential, and vice versa, in all the LTEE lines (Fig. 4A, Table S2). We confirmed two examples of differential gene essentiality in DM25 using clean deletion mutants in the ancestor REL606 and Ara−1 (Fig. S7, Table S1). In total, 77 nonessential genes became essential in at least one evolved lineage, and 97 essential genes became nonessential in at least one evolved lineage, corresponding to ∼17% of the essential genes in the ancestor. Many more genes became nonessential in Ara−6 than in the other evolved lines, as a result of the large duplication discussed below. Across the LTEE populations, we observed a significant tendency for more nonessential genes to become essential than the reverse change (*p* = 0.0071, Mann-Whitney U test). This asymmetry suggests that robustness, in terms of gene essentiality, decreased during the LTEE.

**Fig. 4:**
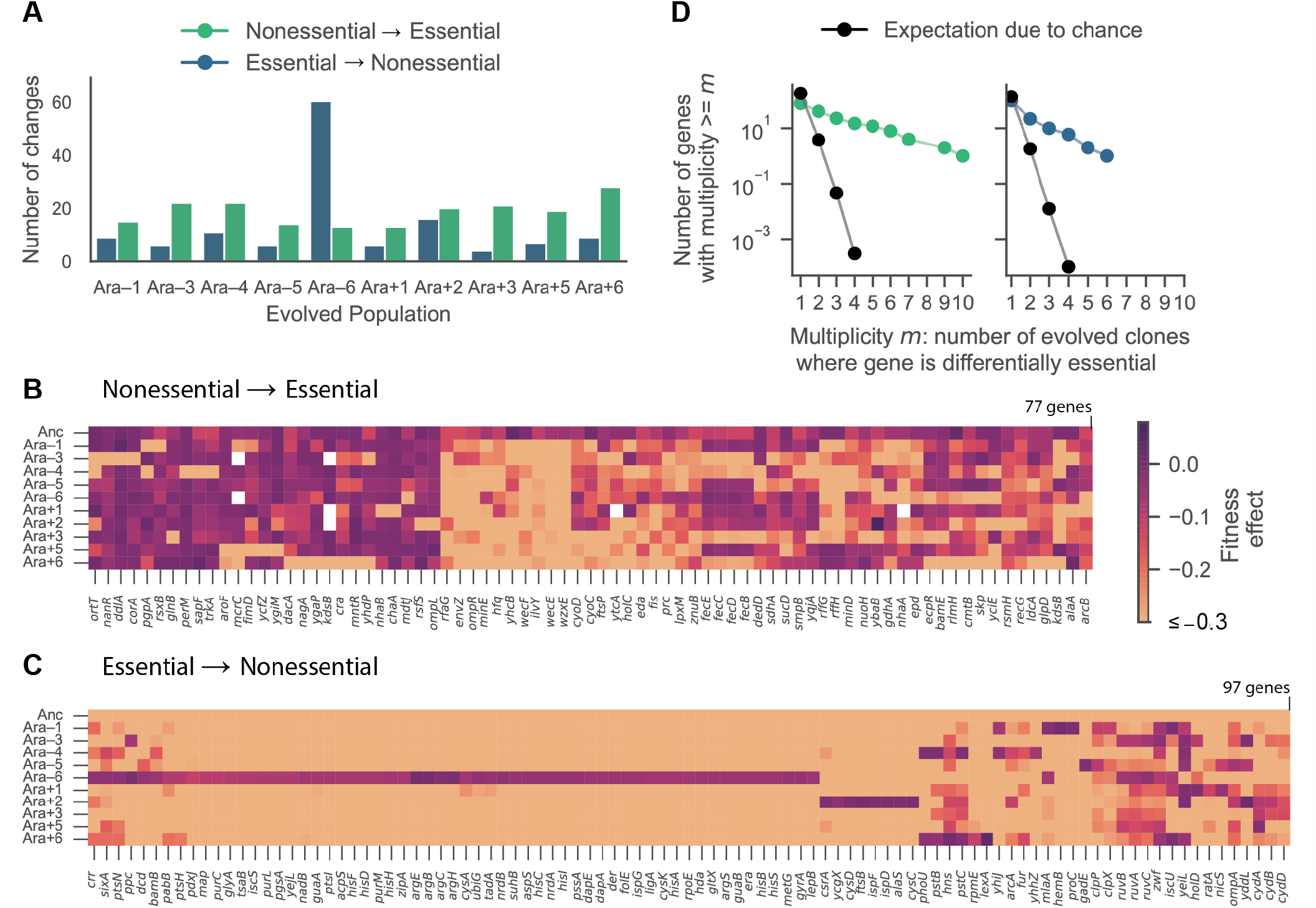
Extensive and parallel changes in gene essentiality over evolution. **(A)** Number of genes that are differentially essential between the ancestor and each evolved strain. **(B, C)** Clustered heatmaps showing fitness effects (scale at right) of genes that evolved to become essential or nonessential in at least one 50,000-generation strain. Genes that were deleted during evolution are shown in white. Genes with mutations conferring fitness effects below –0.3 (the threshold for essentiality) were set to –0.3 for the clustering and visualization. **(D)** Parallel changes in gene essentiality. We estimated the expected number of parallel changes from chance alone by shuffling the gene essentiality and fitness profiles 10,000 times and counting how often the same genes had altered essentially in at least *m* populations. The expectation is an average over 10,000 simulations, and therefore it can be < 1.

Both the essential-to-nonessential and nonessential-to-essential transitions occurred in parallel much more often than expected by chance, similar to the pattern observed for fitness effects (Fig. 3 C, D and E). This outcome was insensitive to the exact cutoff values for essentiality (Fig. S8), and it persisted when we partitioned essentiality changes by the environment (Fig. S9). This parallel evolution strongly implies these changes in essentiality are a result of selection. It is unclear how selection would act directly on essentiality; instead, this parallelism is likely a correlated response of selection for other metabolic or expression changes.

Gene essentiality has previously been associated with highly expressed genes (*30–32*). We therefore examined whether changes in gene essentiality were associated with altered expression levels. We used a recently published RNA-Seq dataset for the LTEE ancestor and evolved strains at 50,000 generations (*33*). Consistent with previous findings, essential genes have higher expression levels on average than nonessential genes (Fig. S10A). However, for those genes that became essential or nonessential during the LTEE, we find no significant differences in the normalized expression levels in the ancestor and evolved strains (Fig. S10B). This result implies that changes in essentiality are not invariably related to altered levels of gene expression.

### Gain or loss of functional redundancy explains some, but not most, changes in essentiality

Gene essentiality may also evolve as by-products of other mutations, such as losses or gains of other gene functions. Structural variation is common in the LTEE populations, with many more large deletions than duplications (*34*). Gene duplications can give rise to robustness because they provide functional redundancy (*35*), whereas deletions might increase the essentiality of other genes by eliminating some redundancies. Because we observed more genes becoming essential than vice versa, we examined whether changes in essentiality might be associated with the presence or absence of redundant genes.

To that end, we sequenced the ancestors and evolved strains at 50,000 generations with high coverage (>60X) and identified all large deletions in the evolved genomes. We identified 251 groups of homologs with >40% identity in the ancestral genome (Table S3). In each evolved strain, between 8 and 36 homolog groups had been reduced to a single surviving member after 50,000 generations.

We therefore asked if the newly essential genes could be explained by the potential loss of functional redundancy caused by these gene deletions. Most evolved strains (all except clones from Ara+5 and Ara+6) have deletions in the *manB-cpsG* region, which spans the *rfb* operon. In the strains with these deletions, insertions in *rffG* and *rffH*, which are paralogs of *rfbA* and *rfbB*, were highly deleterious. The insertion mutants in strains without paralogs had fitness indistinguishable from lethality, whereas the same insertion mutants in strains with the paralogs still present had stable frequencies (Fig. 5A). These results are consistent with a previous finding that the absence of both genes leads to envelope stress owing to accumulation of a toxic intermediate, the ECA Lipid II (*36*). In further agreement, we found that deletion of *wecA*, a gene upstream of ECA II biosynthesis, was tolerated in all strains, indicating that gene is nonessential (Table S4). In another example of gene essentially depending on functional redundancy, a highly expressed copy of *kdsB* became essential owing to either the loss or very low expression of a duplicate copy (Fig. S11, Supplementary Text 2, Table S5). However, such examples were uncommon; for the vast majority of newly essential genes (73 of 77) we found no evidence that essentiality was caused by the loss of redundant genes.

**Fig. 5:**
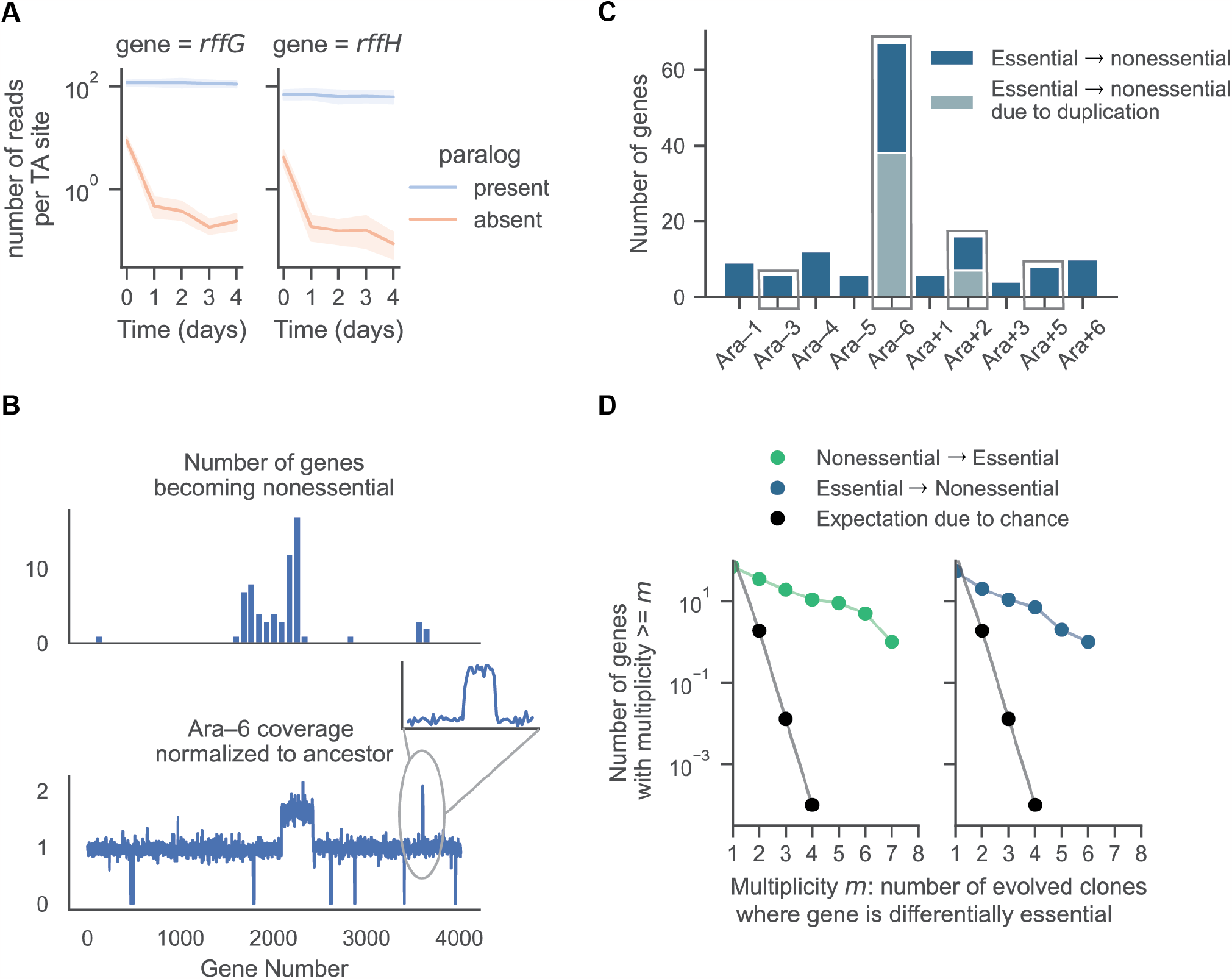
Gain or loss of functional redundancy explains some, but not most, changes in essentiality. **(A)** Loss of paralogs makes *rffG* and *rffH* essential. In strains with *rfbA* and *rfbB* present (ancestors, Ara+5, Ara+6), insertion mutations in *rffG* and *rffH* are nearly neutral, and they persist over the 4-day bulk fitness assay (∼26 generations). In the 8 evolved strains with deletions of the *manB-cpsG* region, insertion mutations in *rffG and rffH* are rapidly outcompeted. Note the logarithmic scale for the abundance; <1 read per TA insertion site is at the level of measurement noise. **(B)** In the 50,000-generation clone from population Ara–6, much of the distribution along the chromosome of the genes that became essential during evolution (top subpanel) maps closely to normalized whole-genome sequencing coverage (bottom subpanel); the inset expands the smaller of two multi-gene duplications. **(C)** Numbers of essential to nonessential changes in the 10 evolved strains. Gray and blue colors represent the numbers that are attributable to gene duplications and those that are not, respectively. Evolved clones with detectable duplications are outlined by a box. **(D)** Parallel changes in gene essentiality, after excluding those changes attributable to the loss or gain of redundancy. We estimated the expected number of parallel changes from chance alone by shuffling the gene essentiality and fitness profiles 10,000 times and counting how often the same genes had altered essentially in at least *m* populations. The expectation is an average over 10,000 simulations, and therefore it can be < 1.

Conversely, we asked whether some genes became nonessential owing to duplication of the region containing them. We first focused on the Ara–6 strain because it had an exceptionally high number of genes that had become nonessential by 50,000 generations. We observed a large duplication in the genome of Ara–6 encompassing ∼300 genes and a smaller duplication spanning ∼25 genes (Fig. 5B). Thirty-eight of the 67 genes that became nonessential in this strain were in those duplicated regions. Three other strains (Ara–3, Ara+2, and Ara+5) had duplicated regions, but only Ara+2 had genes that became nonessential owing to the evolved redundancy (Fig. 5C). Moreover, many genes became nonessential in these and other strains without such duplications. Overall, 52 of 97 genes evolving to become nonessential (∼9% of the ancestral essential genes) were not linked to gene duplications. Thus, structural variation accounts for a little under half the loss of essentiality observed.

Restricting our analysis to changes in gene essentiality not explained by structural variation, we still observed significant parallelism across the independent lineages in genes becoming both essential and nonessential (Fig. 5D). These parallel changes in gene essentiality presumably reflect changes in physiological processes that became either more or less critical for fitness in the LTEE environment.

Some of the consistent changes in essentiality we observed are concordant with prior work on physiological changes in the LTEE populations. In particular, we observed changed essentiality in the Entner-Doudoroff (ED) pathway and in cytokinesis. Metabolic analysis of the LTEE strains identified significantly increased flux through the ED pathway in four evolved strains (Ara+1, Ara+2, Ara+3, Ara–4) (*37*). In this study, we found that disruption of *eda*, an enzyme in the ED pathway, had highly deleterious effects in those strains, suggesting the ED pathway plays an important role in increased fitness in the LTEE. Cell size has increased substantially over the LTEE (*38*), potentially reducing the nucleoid/volume ratio. Our results show that genes involved in cytokinesis (*minD, minE, dedD, ftsP*) became more essential in the evolved populations. We hypothesize that, as cells become larger, disrupting the min system became more deleterious as the nucleoid occupies relatively less volume. As a consequence, the nucleoid occlusion system may become less effective at preventing aberrant Z-ring formation, thereby increasing the frequency of minicell formation and reducing fitness. More generally, we hypothesize that many other as yet unexplained changes in gene essentiality are similarly the indirect result of selection on various metabolic pathways.

## Discussion

Biological systems, from proteins and genetic networks to organismal physiology and fitness, are often remarkably robust to mutations, but how robustness evolves is not well understood (*12, 39–41*). Here, we show that overall robustness to mutations, measured as the average fitness effect of insertion mutations in bacterial genomes, did not change systematically during 50,000 generations of the LTEE, despite significant changes in the fitness effects of mutations in many genes. However, for a subset of cases, we observed a bias towards more genes becoming essential than nonessential. This asymmetry lends some support to the “increasing costs” model of epistasis for mutations in those genes subject to the largest shifts in fitness effects over evolution. This pattern is thus consistent with previous findings in yeast of a tradeoff between fitness and robustness in large-effect mutations (*17*), but this effect disappears when looking globally. Overall, our results paint a complex picture of changing fitness effects that no simple model adequately captures.

Despite the lack of consistent directional changes in the overall statistical properties of the DFEs, a key finding of our study is that the set of essential genes changed often during the LTEE, with many nonessential genes becoming essential and many essential genes becoming nonessential. While prior work has shown that gene essentiality is not a static property of a species, but instead is evolvable (*18, 22, 23*), our results show that gene essentiality changes consistently, even without direct selection for suppression of essentiality.

The 125 genes (∼3% of the genome) that changed in essentiality over 50,000 generations in a single environment is comparable to the 120 variably essential genes between diverse strains of *E. coli* when tested across three environment conditions (*20*) and separated by vastly more generations. The same study also suggested that horizontal gene transfer (HGT) has played a major role in driving changes in gene essentiality in bacteria (*20*). However, the ancestral *E. coli* strain used in the LTEE lacks plasmids, functional prophages, and natural transformation (*42*). Thus, our results show that gene essentiality can evolve rapidly even without HGT.

The ability to predict evolution requires a deep understanding of fitness landscapes and how they change. Our study shows that, while the overall distribution of fitness effects was largely unchanged during the LTEE, many individual mutations became more or less deleterious owing to epistatic interactions with the evolving genetic background. As a consequence, evolutionary paths that were inaccessible to the ancestor have become available, while others are closed off, as recently described in the context of protein evolution (*43*). While natural selection steers replicate populations along similar trajectories (*44–46*), the paths that open or close are often the same across independently evolving lineages. Taken together, these results demonstrate the dynamic nature of gene essentiality, and more broadly, the fitness effects of mutations, and they show that access to some evolutionary trajectories changes consistently over time, even as the macroscopic features of the fitness landscape remain largely unchanged.

## Supporting information

Supplemental methods, figures, and protocols

## Acknowledgments

We thank Milo Johnson, Michael Desai, Andrew Murray, Thomas Bernhardt, Alejandro Couce, Olivier Tenaillon, Célia Souque, Fernando Rossine, and Daniel Eaton for valuable feedback and discussion; Thao Truong, Joel Sher, and Thomas Bernhardt for sharing *E. coli* MFDpir and donor plasmid pSC189, and for assistance with generating transposon libraries; Tanush Jagdish for help with the LTEE strains and culture conditions; and Karel Brinda and Natalia Qui ñones-Olvera for assistance with the bioinformatics analyses.

## Funding

A.L. acknowledges support from the Molecules, Cells, and Organisms Graduate Program, Harvard University. R.E.L. acknowledges support from the US National Science Foundation (DEB-1951307) and the John Hannah endowment at Michigan State University. M.B. acknowledges support from the NIGMS of the National Institutes of Health (R35GM133700), the David and Lucile Packard Foundation, the Pew Charitable Trusts, and the Alfred P. Sloan Foundation. Sequencing was performed at the Bauer Core Facility at Harvard University, and computational work used the O2 cluster supported by the Research Computing Group at Harvard Medical School.

## Author Contributions

A.L. and M.B. designed the project; A.L., S.V.O. conducted the experiments, A.L. generated the sequencing data; A.L., S.V.O. and M.B. designed and troubleshot the experiments; A.L. designed and conducted the bioinformatics analyses; A.L., C.M.H., R.E.L., and M.B. designed statistical analyses; A.L. analyzed the data; R.E.L. directs the LTEE and provided strains and critical feedback on interpretation; all authors wrote and revised the manuscript.

## Competing interests

The authors declare no competing interests.

## Data Availability

Raw sequencing reads have been deposited in the NCBI BioProject database under accession number <u>PRJNA814281</u>. Processed data were deposited on Zenodo (https://doi.org/10.5281/zenodo.6547536); source code for the sequencing pipeline, downstream analyses, and figure generation are at GitHub (https://github.com/baymlab/2022_Limdi-TnSeq-LTEE).

## Supplementary Material

Materials and Methods

Supplementary Text

Figs. S1 to S11

List of Tables S1 to S5

Detailed Experimental Protocols

References (47-55)

## Notes

https://github.com/baymlab/2022_Limdi-TnSeq-LTEE

https://doi.org/10.5281/zenodo.6547536

### Competing Interest Statement

The authors have declared no competing interest.

### Summary of Updates

Manuscript updated and revised, particularly title and introduction, to reflect broader scope of results

