## Supplemental methods, figures, and protocols for "Parallel evolution of mutational fitness effects over 50,000 generations"

**This PDF file includes:**

Materials and Methods
Supplementary Text
Figures S1 to S11
List of Tables S1 to S5
Detailed Experimental Protocols
References

### Materials and Methods

#### *Strains, Plasmids and Growth Conditions*

We used the LTEE ancestors (*E. coli* strains REL606 and REL607, called ANC and ANC*, respectively, in our paper) and evolved clones (listed below) sampled from all 12 populations at 50,000 generations.

| Population | Clone ID | Population | Clone ID |
|---|---|---|---|
| Ara–1 | REL 11330 | Ara+1 | REL 11392 |
| Ara–2 | REL 11333 | Ara+2 | REL 11342 |
| Ara–3 | REL 11364 | Ara+3 | REL 11345 |
| Ara–4 | REL 11336 | Ara+4 | REL 11348 |
| Ara–5 | REL 11339 | Ara+5 | REL 11367 |
| Ara–6 | REL 11389 | Ara+6 | REL 11370 |

We generated the donor strain for transposon library construction by transforming *E. coli* MFDpir with pSC189, a mobilizable plasmid that carries the mariner transposon but lacks the machinery necessary for replication outside the MFDpir host. We grew the donor strains in LB with 300 μM DAP (diaminopimelic acid) and 50 mg/L kanamycin. We produced a fresh batch of donor strains each week, to ensure that the transposon had not been mobilized or disrupted essential genes in the donor background. Single gene deletions were constructed by recombineering, as previously described (*47*). We used the pSIM5-tet plasmid carrying a heat-inducible lambda red recombinase and a tetracycline-resistance gene (*48*). The kanamycin-resistance cassette used for recombineering was amplified from the pKD4 plasmid (*47*).

#### *Transposon Library Construction*

We used a suicide-plasmid delivery system to construct the transposon libraries (*49*, *50*). We conjugated dense overnight cultures of the *E. coli* MFDpir-pSC189 donor and recipient (one of the LTEE strains) on a 0.2-μm filter and incubated for 1 h. We resuspended the conjugation mixture and plated 1-2 mL on large LB agar plates supplemented with kanamycin (either 50 mg/L or 100 mg/L, depending on selection efficiency). After overnight growth at 30°C, we scraped cells from the plates, mixed the resulting mutant library with glycerol, and stored the library at –80°C. We plated dilutions of the transposon libraries and verified that they carried kanamycin resistance by patching ~50 colonies on LB and LB + kanamycin plates. We only retained those libraries where >90% colonies were kanamycin resistant.

#### *Fitness Assays*

We performed bulk fitness assays in DM25 (Davis-Mingioli minimal medium with 25 mg/L glucose) held in glass test tubes and incubated at 37°C with 220 rpm orbital shaking. We used test tubes, instead of the small Erlenmeyer flasks used in the LTEE, in order to run many fitness assays simultaneously. Previous work found no systematic difference in fitness estimates using glass tubes versus flasks.

Each fitness assay comprised five 10-mL cultures propagated in parallel. This replication increased the bottleneck population size by a factor of five, without introducing any density-dependent effects. For each assay, we inoculated 50 mL of DM25 with ~$5 \times 10^5$ cells/mL from the freezer stock of the transposon library (comparable to the number of cells transferred during the LTEE), and we then split the volume among five glass tubes. After incubating for 24 h at 37°C with shaking, we pooled the five tubes. For four days (~26.5 generations in total) we diluted 500 μL of each bulk competition into 50 mL fresh DM25 (1:100 daily dilution, as in the LTEE) and immediately split the volume among five cultures, as before. We spun down the remaining cells, and we stored the pellet frozen for later DNA extraction and analyses. For each transposon library, we performed two replicate fitness assays, starting from the same aliquot of the transposon library from the freezer stock (which served as the time-zero point).

***Recombineering and Pairwise Fitness Assays***

We grew overnight cultures of the strains for recombineering (containing the pSIM5-tet plasmid) in LB at 30°C, and we diluted them 100-fold in 50 mL of low-salt LB. When the cultures were in mid-exponential phase (OD ~0.4), we transferred them to a water bath at 42°C to induce the recombinase for 15 min. We then chilled the cultures on ice, and we prepared competent cells from the induced cultures by washing and pelleting cells multiple times in 10% glycerol. We then transformed the competent cells with 200-500 ng of the recombineering insert, which had 40 bp identical to the flanking regions of the gene we sought to delete as well as the kanamycin-resistance cassette (primer sequences shown in Part 6 of Detailed Experimental Protocols). The recombineering protocol worked successfully for the LTEE ancestor (REL606), but we were unable to make deletion mutations in the evolved strains except the 50,000-generation clone from population Ara–1 (REL11330), for which we saw a reduced efficiency. We found that the limiting factor for most evolved strains was transformation of the recombineering plasmid into the cell using electroporation.

For pairwise competitions, we grew overnight cultures of the unmutated parent and the deletion mutant in LB. We mixed them 1:1 volumetrically and inoculated the mix in 10 mL of DM25 with ~$5 \times 10^5$ cells/mL. We spread dilutions from the competition mixture on LB and LB + Kanamycin plates at the start of the competition and after one day in DM25. The counts on the LB + Kanamycin plates correspond to the number of mutant cells, while the counts on the LB plates correspond to the total number of cells including both the parents and mutants. We estimated the fitness effect of the deletion mutation as the rate of change in the ratio of the mutants to the parents, as follows:

$$s = \ln \frac{N_{LB+Kan,t1} \,/\, (N_{LB,t1} - N_{LB+Kan,t1})}{N_{LB+Kan,t0} \,/\, (N_{LB,t\,0} - N_{LB+Kan,t0})} \,/\, \log_2(100)$$

where $\log_2(100)$ is the number of generations (doublings) per day during the competition given the 100-fold dilutions.

***DNA Extraction, UMI-TnSeq Library Preparation, and Sequencing***

We used the Invitrogen PureLink gDNA extraction kit to extract DNA from ~$2 \times 10^9$ cells for each transposon library. We measured DNA concentrations using the Invitrogen Quant-it Kit and normalized them to 20 ng/μL. We ran 10 μL tagmentation reactions with 5 μL Illumina TDE1 buffer, 2.5 μL TDE1 enzyme, and 2.5 μL normalized DNA at 55°C for 10 min. We used the entire volume of the reaction as the template for PCR1, a low-amplification cycle PCR where we added unique molecular identifiers using custom primers (see Part 4 of Detailed Experimental Protocols for primer sequences). We cleaned the product of PCR1 with 1.2X serapure beads, which we eluted in 15 µL $dH_20$. We used the eluant as template for PCR2, where we selectively amplified fragments containing the transposon sequence. We cleaned the product of PCR2 with 1.2X serapure beads, which we eluted in 25 µL $dH_20$. We quantified the concentration of DNA of the amplified transposon libraries using the Invitrogen Quant-it Kit, and we diluted and pooled all samples to 4 nM. We verified the concentration of the pooled library using the Kapa Biosystems Illumina qPCR kit. We also prepared whole-genome sequencing libraries for all the strains using the tagmentation based-approach from Baym et al (*51*). Additional details on the UMI-TnSeq library preparation are included in Part 3 of the Detailed Experimental Protocols. We sequenced the transposon and whole-genome libraries on two lanes of an Illumina NovaSeq S4 (paired end, 2 x 150 bp) at the Bauer Core Facility, Harvard University.

***Data Analysis: Fitness and Essentiality***

We obtained demultiplexed reads from the sequencing core. We filtered reads using a custom Python script, retaining only those reads that contained a sequence matching the end of the mariner transposon, and we stored the 10-bp unique molecular identifier (UMI) sequence separately. We used bowtie2 to map the filtered reads to the reference genome of the LTEE ancestor, *E. coli* strain REL606. We extracted coordinates of all the uniquely mapped reads; these coordinates correspond to the TA sites in the reference genome. For every TA insertion site, we identified corresponding UMIs and mapping coordinates, and we used this combination to count the number of distinct biological replicates for each TA site. We obtained a median of 23 million reads per timepoint in the fitness assays. Approximately 85% of the reads mapped to the reference genome; ~20% of those reads were discarded as PCR duplicates. We consolidated all the data from the bulk fitness assay into a master file for each transposon library; that dataset includes the counts for every insertion site at each timepoint in the fitness assay.

For the downstream analyses, we normalized each insertion by the total sample size. For each insertion mutation with at least 5X coverage at the start of the assay, we estimated its fitness as the slope of the linear fit to its ln(frequency) over time (in number of generations, $\log_2(100) \cong 6.7$ per day). As explained elsewhere, this metric differs from the ratio of Malthusian growth rates (as used in many other LTEE analyses) by a factor of ln 2 (*52*). For those mutants that disappeared, whether due to chance or the mutant being deleterious, we added a pseudo-count of 1. For each gene, we then averaged over all the insertion sites using an inverse-variance weighting approach, excluding the first 10% and last 25% of the gene's length.

We estimated the error in the fitness measurement of a gene as the weighted standard error of the mean of the fitness estimates for all *K* interior TA sites within the gene:

$$\mu_{gene} = \left(\frac{1}{K}\right) \sum_{i \in TA\ sites} w_i s_i.$$

The weights were defined as follows:

$$w_i = \frac{log(2)^2}{\left(\frac{1}{1+n_0} + \frac{1}{1+n_1}\right)},$$

where $n_0$ and $n_1$ are the number of reads at timepoints 0 and 1, respectively, and $K$ is the number of TA sites. Maximum weights were set with $n_0$ and $n_1$ both equal to 100. We define the error in the fitness estimate of a gene as the inverse-variance weighted standard error of the mean:

$$\Delta\mu_{gene} = \sqrt{\frac{1}{(N)(N-1)}(\sum_{i\in TA\,sites} w_i(s_i - \mu_{gene})^2} \ .$$

The fitness effect of each gene was adjusted by a correction factor equal to the average fitness effect of insertions that disrupt pseudogenes. This adjustment ensures that our fitness estimates are not influenced by changes in the mean population fitness during the bulk fitness assay. The mean of the DFE is:

$$\mu_{DFE} = \sum_{genes}\mu_{gene} \,/\, N - \sum_{pseudogenes}\mu_{pseudogene} \,/\, M \ ,$$

where $N$ and $M$ are the numbers of genes and pseudogenes, respectively. We define the uncertainty in the estimate of the mean of the DFE as:

$$\Delta\mu_{DFE} = \sqrt{\sum_{genes}(\Delta\mu_{gene}\frac{\partial\mu_{DFE}}{\partial\mu_{gene}})^2 + \sum_{pseudogenes}(\Delta\mu_{pseudogene}\frac{\partial\mu_{DFE}}{\partial\mu_{pseudogene}})^2} \text{ , where}$$

$$\frac{\partial\mu_{DFE}}{\partial\mu_{gene}} = 1/N, \frac{\partial\mu_{DFE}}{\partial\mu_{pseudogene}} = 1/M \ .$$

We restricted our analysis to those protein-coding genes with at least 5 interior TA sites; we further required that a gene had at least two trajectories meeting thresholds in both technical replicates, and that at least 20% of the gene's TA sites were used in the fitness estimation. We chose these thresholds to ensure that we had sufficient data for every gene that was included in our analyses of changes in essentiality, and to exclude potential outlier sites in essential genes from contributing to the fitness estimates. We realize that excluding 10% and 25%, respectively, of the 5' and 3' ends of a gene is an imperfect solution, but it greatly reduced the impact of outliers. We also excluded all genes with annotations linked to transposons or mobile genetic elements, because they might move between sites in the genome. When comparing fitness estimates between two strains, we often found that fitness effects were calculated in one strain but not the other owing to these arbitrary thresholds. To get around this problem, we calculated fitness effects in such cases using relaxed thresholds (i.e., at least one trajectory meeting thresholds in both replicates, instead of two as otherwise required).

To classify a gene as differentially essential between ancestral and evolved strains in DM25, we required mutations in that gene to have a fitness effect > –0.15 in one strain and < –0.3 in the other. The inclusion of the –0.15 threshold was to ensure that the difference in fitness effects between strains is sufficiently large that it cannot be attributed to measurement noise. The choice of the –0.3 threshold is based on numerical simulations of fitness effects for genes that are essential for growth in DM25, which we used to identify the most deleterious fitness effect such that the overlap in the distributions for an essential gene and a non-essential gene subject to the merely deleterious mutations is < 0.05 (Fig. S6).

The simulations were performed as follows:

- Fit a distribution to the number of reads per TA site from experimental data using the ANC strain at the initial timepoint $t_0$ (restrict to sites with at least 10 reads). We fit both normal and gamma distributions to the log-transformed number of reads per TA site. Both distributions work well, but we report results using the gamma distribution.

- Draw an initial abundance from the fit to experimental data, which is the number of reads at $t_0$ and called $N_0$.
- Estimate the expected number of reads at $t_1$ as follows:
    - $E[N_1] = N_0/100 * \exp((1 + s)6.64)) = N_0 * \exp(6.64s)$ if the gene is not essential. The value 6.64 equals the number of generations (doublings) of population growth per day, which offsets the 100-fold daily dilution.
    - $E[N_1] = N_0/100$, if the gene is essential, such that the mutant does not grow at all. The division by 100 reflects the daily 100-fold dilution without growth.
    - Draw from a Poisson distribution with mean $E[N_1]$, which we call $N_1$.
- Estimate the simulated fitness effect as:
    - $s = \ln((N_1 + 1)/N_0 + 1))/6.64$.
- Repeat the simulation 5,000 times.

Even with relaxed thresholds for fitness estimation, there were many cases of genes that were nonessential in DM25 in one strain but did not have any estimated fitness in the other strain. To determine whether these cases arose because insertion sites were missing by chance, or whether the gene was differentially essential in LB, we used the following approach:

- We first define essentiality in LB based on the fraction of TA sites, $f$, in a gene that had at least 5 reads mapping to a site at time 0 (after correcting for sequencing depth). We classified those genes with $f < 0.1$ as essential in LB (Fig. S6A).
- For the LTEE ancestor REL606, this definition captured 88% of the essential genes in *E. coli* K-12 (*25*), with a false positive rate of only 2%. The area of the receiver-operator characteristic curve is 0.96 (Fig. S6B), which validates our approach to defining a gene as essential in LB.
- Next, identify all genes that are nonessential ($s > -0.15$) in DM25 in strain 1, and with fitness not calculated in strain 2, with fraction $f < 0.1$.
- For every such gene, calculate the number of TA sites at time 0 with coverage > 5X in both strain 1 ($n_1$) and strain 2 ($n_2$).
- Calculate the probability of observing $n_2$ or fewer sites, assuming that the expected number of sites is $n_1$.
- Adjust those $p$-values using the Benjamini-Hochberg False Discovery Rate (FDR).
- Define those genes for which this FDR-adjusted probability is < 0.05 as differentially essential in LB for the two strains in question.

This approach ensures that we are conservative in calling genes differentially essential, particularly in those genes with fewer than 10 insertion sites. As an additional control, using the same parameters as described above, we did not find any genes that are differentially essential between the two marked variants of the LTEE ancestors (strains REL606 and REL607, which we call Anc and Anc*, respectively) that have repeatedly been shown to have equal fitness.

**Estimating the number of essential genes in the LTEE ancestor in DM25**

We counted the number of genes where disruptions led to a fitness estimate of $s < -0.3$. To account for the fact that some genes may be missing from the initial transposon library by chance (i.e., due to very few TA sites within the gene's interior), we excluded those genes with an FDR-corrected probability > 0.05 (as described for defining differential essentiality in LB).

***Data Analysis: Structural Variation***

The average depth of coverage of the whole-genome sequencing data was > 60X. We used breseq (*53*) to identify regions of the genome that had been deleted in the evolved strains during the LTEE. We identified deletions > 1 kb using the missing-coverage evidence from the breseq output files. To identify duplicated regions, we used the samtools depth command (*54*) to get depth of coverage at every site in the genome, using the ancestral REL606 genome as the reference. To account for variability in coverage across the genome, we calculated the normalized coverage for each evolved strain relative to REL606. We obtained the coordinates for duplicated genes by visualizing the normalized coverage and identifying those regions where the coverage was consistently 1.5X or higher than the background level. We set the cutoff at >1.5X coverage over background because large duplications also have a tendency to be lost when clones are being grown, as is done to obtain their DNA for sequencing.

To identify gene homologs, we used MMseqs2 cluster (with parameters: --min-seq-id 0.4) with a 40% identity threshold for homologs (*55*). For each evolved clone, we asked whether all but one member of the paralogous group had been lost during evolution and, if so, we evaluated whether the other member had become essential.

### Supplementary Text

1. Exclusion of Ara–2 from further analysis

We excluded the Ara–2 sample from all comparisons after we found that fitness measurements in this clone were unreliable and systematically biased. In this genetic background, a few insertion mutations massively increased in frequency during the fitness assay, depressing the frequencies of other mutations. In principle, we might compensate for this effect by excluding the outliers, and then correcting the other mutations using pseudogenes as the neutral expectation. However, the insertion mutations in pseudogenes were also outcompeted in the fitness assay; their counts fell to nearly zero on day 4, and the estimated fitness effects for these insertions range from –0.2 to –0.15. As a consequence, there are two unavoidable problems with using the pseudogenes to establish a correction factor for the remainder of the DFE:

- Measurements of deleterious fitness effects are inherently noisier than nearly neutral effects, leading to generally larger errors and potentially systematic biases for fitness estimates of all other mutations if we used the mean pseudogene value as a correction factor.
- By subtracting a large, constant deleterious fitness effect (nearly –0.2) from all other measurements, the distribution of fitness effects in Ara–2 would abruptly end at $s \cong -0.3$. This approach would effectively truncate the deleterious tail of the DFE, which in other backgrounds spans effects as deleterious as –0.6 to –0.3. Consequently, a calculated fitness effect of –0.25, for example, could not be meaningfully compared between Ara–2 and other populations.

2. Expression Levels and *kdsB* Gene Essentiality

The gene *kdsB* is essential in *E. coli* strain K-12. However, there are two copies of *kdsB* in the LTEE ancestor (ANC, REL606), and both copies have high expression levels (*33*), making each copy dispensable. In three evolved populations (Ara–3, Ara+1, Ara+2), one copy of *kdsB* (indicated as *kdsB_2* in Table S5) has been lost, making the other copy (*kdsB_1*) essential. In strains with two copies of *kdsB*, whether *kdsB_1* is essential or not depends on the expression level of *kdsB_2*. The fitness effect of insertions in *kdsB_1* is correlated with the expression level of *kdsB_2* (Fig. S11); in strains with high expression levels of *kdsB_2*, the effect of disrupting *kdsB_1* is less deleterious, and vice versa. This example shows that the presence of a duplicate gene is not sufficient to make a gene dispensable; instead, the duplicate must also be expressed at a high enough level.

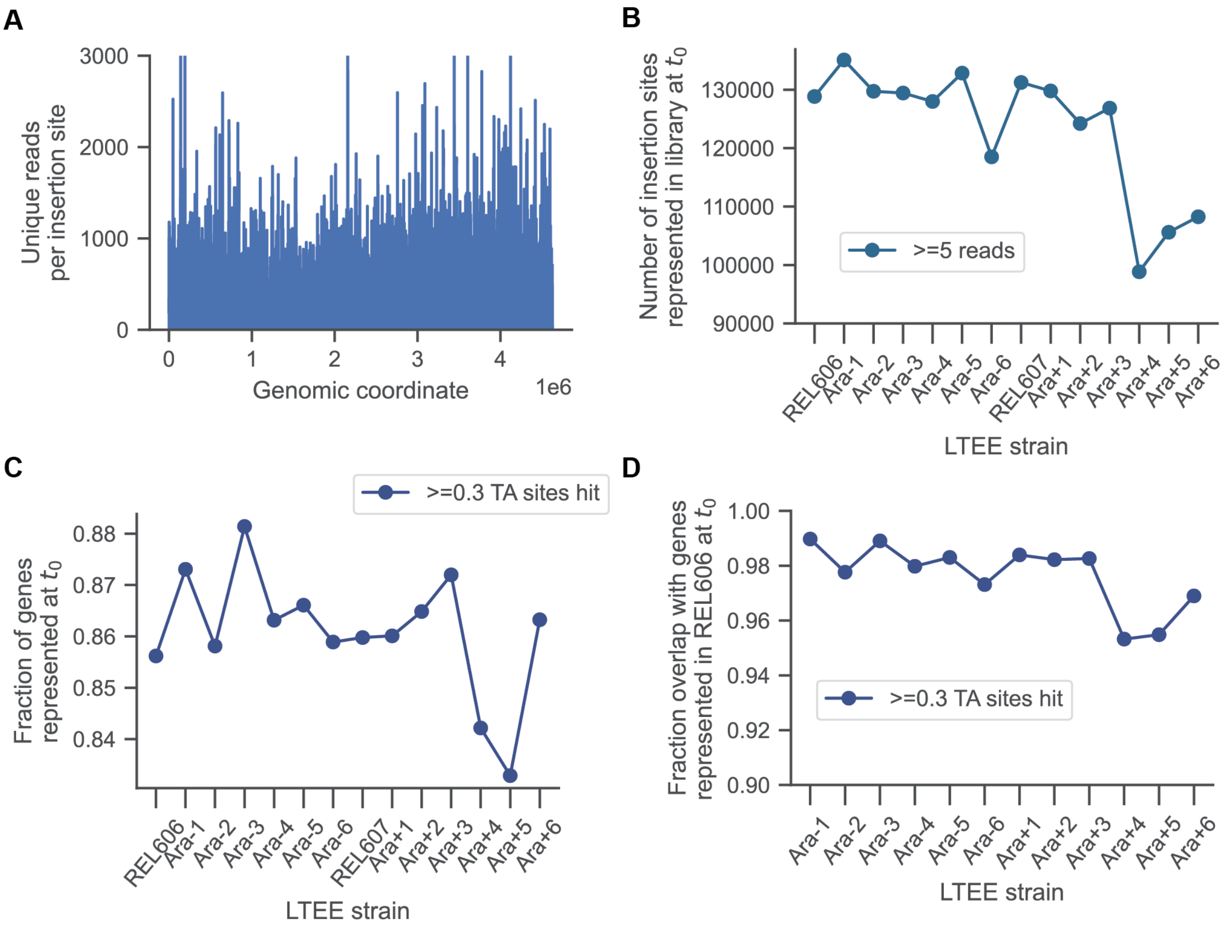

**Fig. S1: Distribution of transposon insertions and representation of genes in the transposon sequencing dataset. (A)** Insertions in the ancestor REL606 (a representative example) are distributed throughout the genome. **(B)** Number of insertions at the timepoint zero with at least five mapped reads (Note that the total number of available insertions in the ancestor is 211,995). **(C)** Fraction of genes represented (at least 30% of the interior TA sites with 5 or more mapped reads) at the timepoint zero. **(D)** Fraction of genes represented in the ancestor, REL606, that are also represented in the evolved strains.

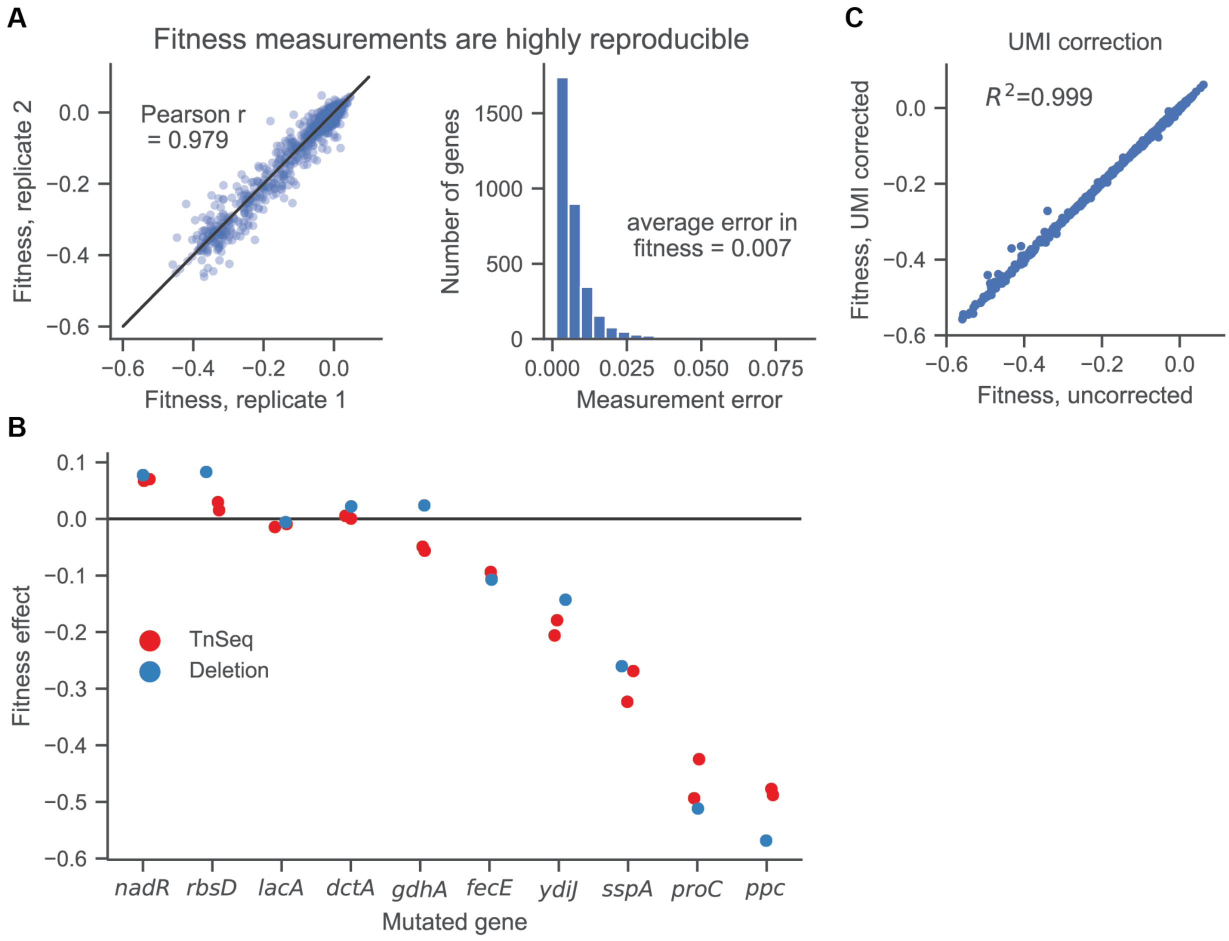

**Fig. S2: Estimates of fitness effects of TnSeq insertion mutations are highly reproducible and consistent with fitness effects of deletions in the same genes**. **(A)** Correlation between technical replicates in evolved Ara+1 clone (left) and distribution of measurement errors (right). **(B)** The fitness effects estimated from disrupting genes using TnSeq in the LTEE ancestor and by pairwise competitions between the unmutated ancestor and clean deletion mutations are consistent. Each point represents an independent measurement. **(C)** Fitness estimates with and without UMI correction are very consistent with each other.

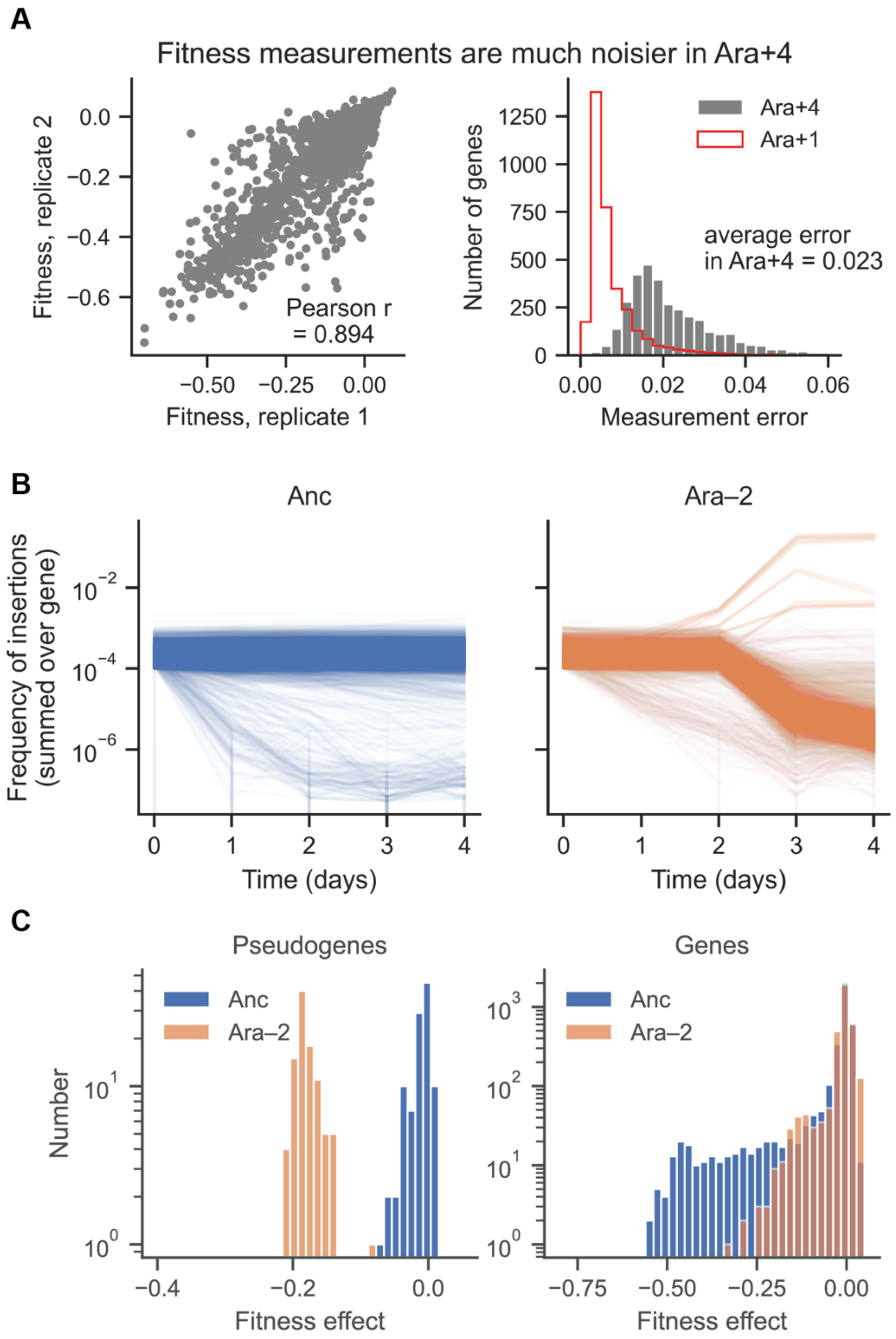

**Fig. S3: Evolved strains from populations Ara–2 and Ara+4 were excluded from further analyses because of aberrant properties. (A)** The correlation between technical replicates is much worse for Ara+4, and the typical measurement error much larger (2.3%), than for other strains (ranges from 0.7 to 1.2%). **(B)** Plot of all mutant trajectories in the LTEE Ancestor and Ara–2. **(C)** Fitness estimates are biased in Ara–2 because of the highly negative correction factors based on the pseudogenes (left panel). As a consequence, the DFE for Ara–2 barely extends below –0.3 (right panel).

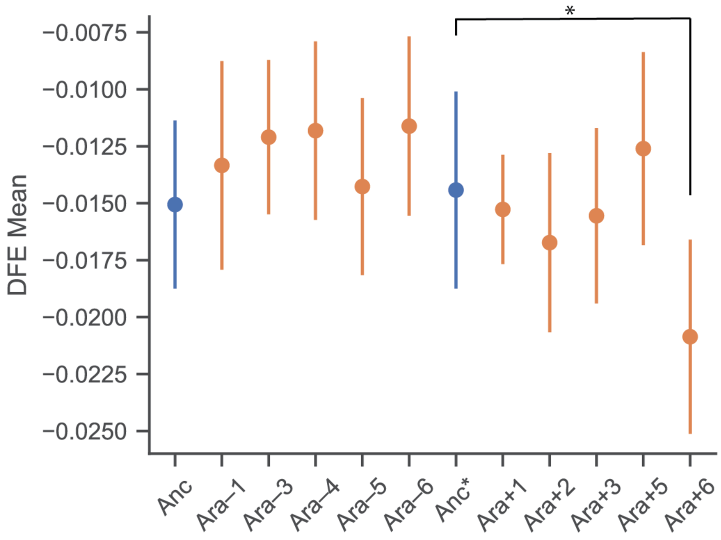

**Fig. S4: Differences in the DFE are largely driven by the deleterious tail of fitness effects** ($Z$-test, $*$ $0.01 < p < 0.05$). Here, we compared the means of the DFE for the populations excluding mutations with a fitness effect $s < -0.3$, as these measurements are noisier, and prone to biases due to differences in number of surviving cells, and therefore generations of growth after recovery from the freezer stock.

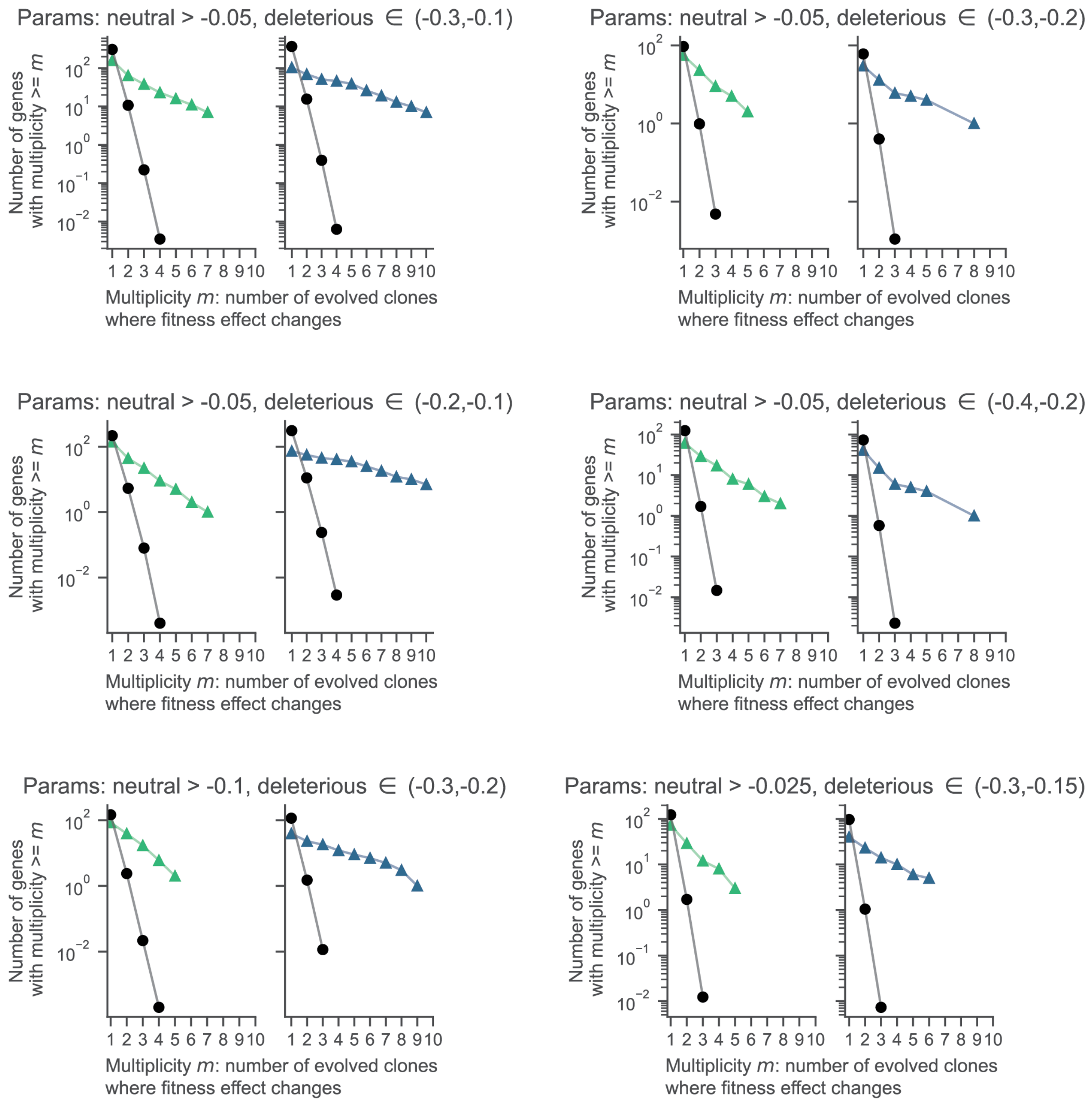

815 **Fig. S5: Signature of parallelism in fitness effect changes is robust to the choice of thresholds for defining neutral and deleterious fitness effects.** The number of genes with varying fitness vastly exceeds what would be expected from pure randomness in all these plots. The expectation is an average over 10,000 simulations, and therefore it can be < 1.

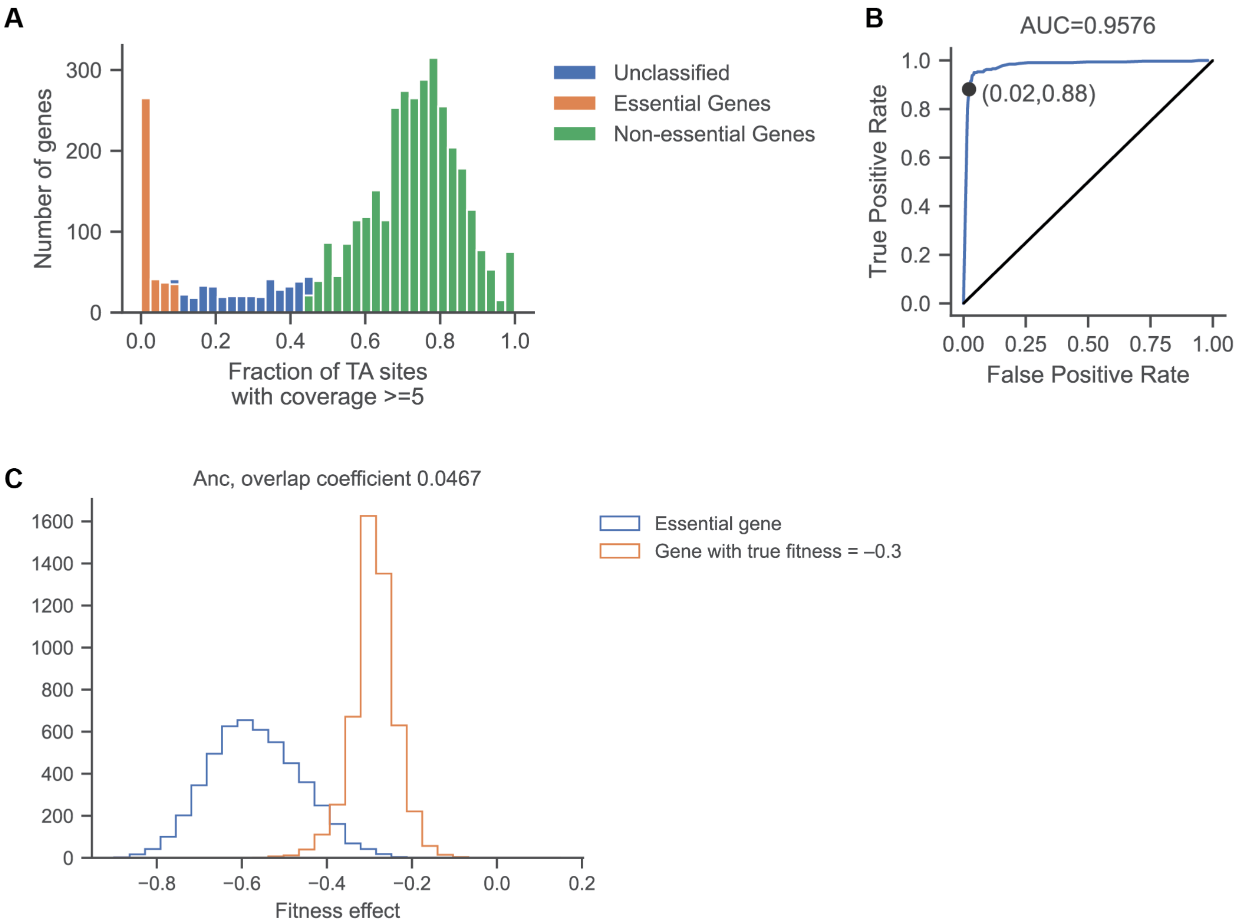

**Fig. S6: Defining gene essentiality in LB and DM25. (A)** LB: Distribution of fraction of sites with coverage $\geq 5$ at $t_0$ in the fitness assay in the ancestor, REL606. Genes with that fraction $< 0.1$ are classified as essential and genes with fraction $> 0.45$ as nonessential. **(B)** Classification of gene essentiality in LB is reliable when compared to data from Goodall et al. (*25*). **(C)** DM25: We simulated the fitness estimates for a truly essential gene (no growth during the assay) and for genes with true fitness effects ranging from –0.45 to –0.2. We found the most deleterious effect, $s = -0.3$, where the overlap coefficient is $< 0.05$. This overlap represents the probability of observing the same fitness effect in a truly essential gene and in a gene disruption that is highly deleterious, and therefore the limit of our fitness assay to distinguish essentiality from significant growth deficit. See Methods: *Data Analysis: Fitness and Essentiality* for a detailed description of these simulations.

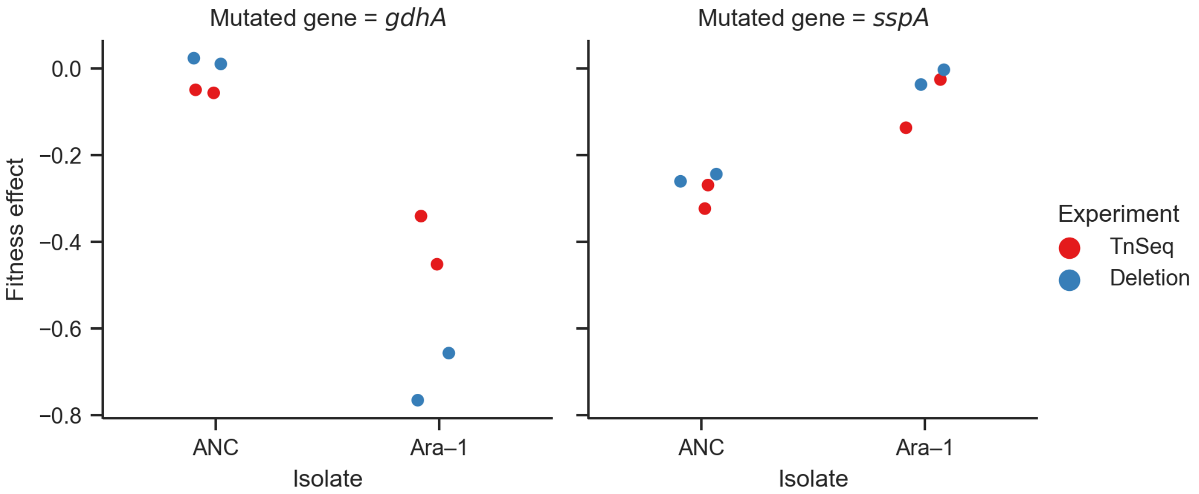

**Fig. S7: Verification of differential gene essentiality for two genes (*gdhA* and *sspA*) in the ancestor and an evolved clone from population Ara–1, obtained by generating clean gene deletion mutants in both genetic backgrounds.** For both genes, the prediction of differential essentiality from TnSeq is consistent with competition assays using the deletion mutants. For *gdhA* in the Ara–1 background, both methods classify the gene as essential ($s < -0.3$). The difference between the two methods can be large for essential genes, because the measurement errors for both methods are very large as a consequence of the rapid disappearance of unfit mutants that results in extremely low counts.

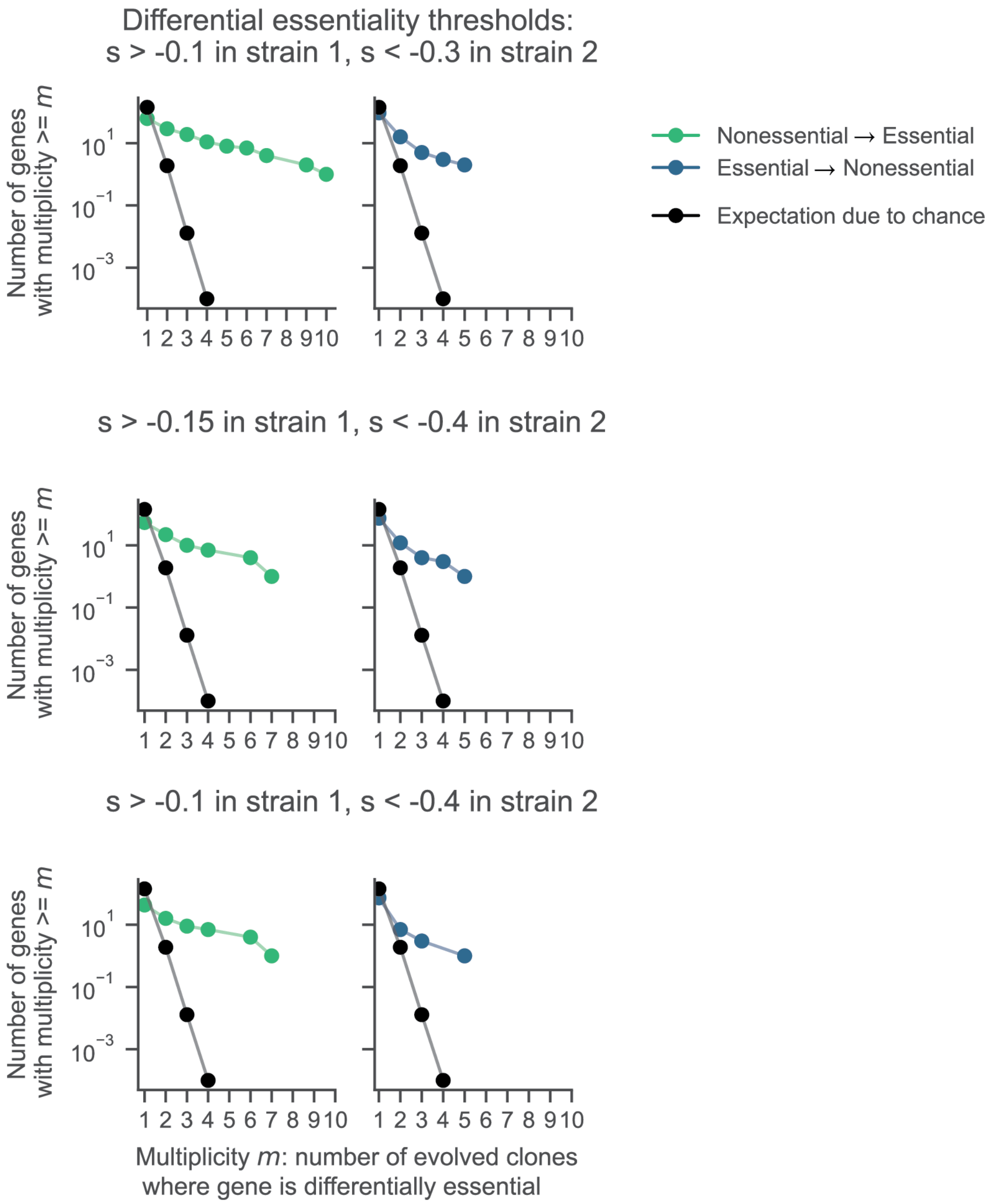

**Fig. S8: Signature of parallelism in changing gene essentiality is robust to the choice of thresholds for defining differential essentiality.** The number of genes with changed essentiality vastly exceeds what would be expected from pure randomness in all these plots. The expectation is an average over 10,000 simulations, and therefore it can be < 1.

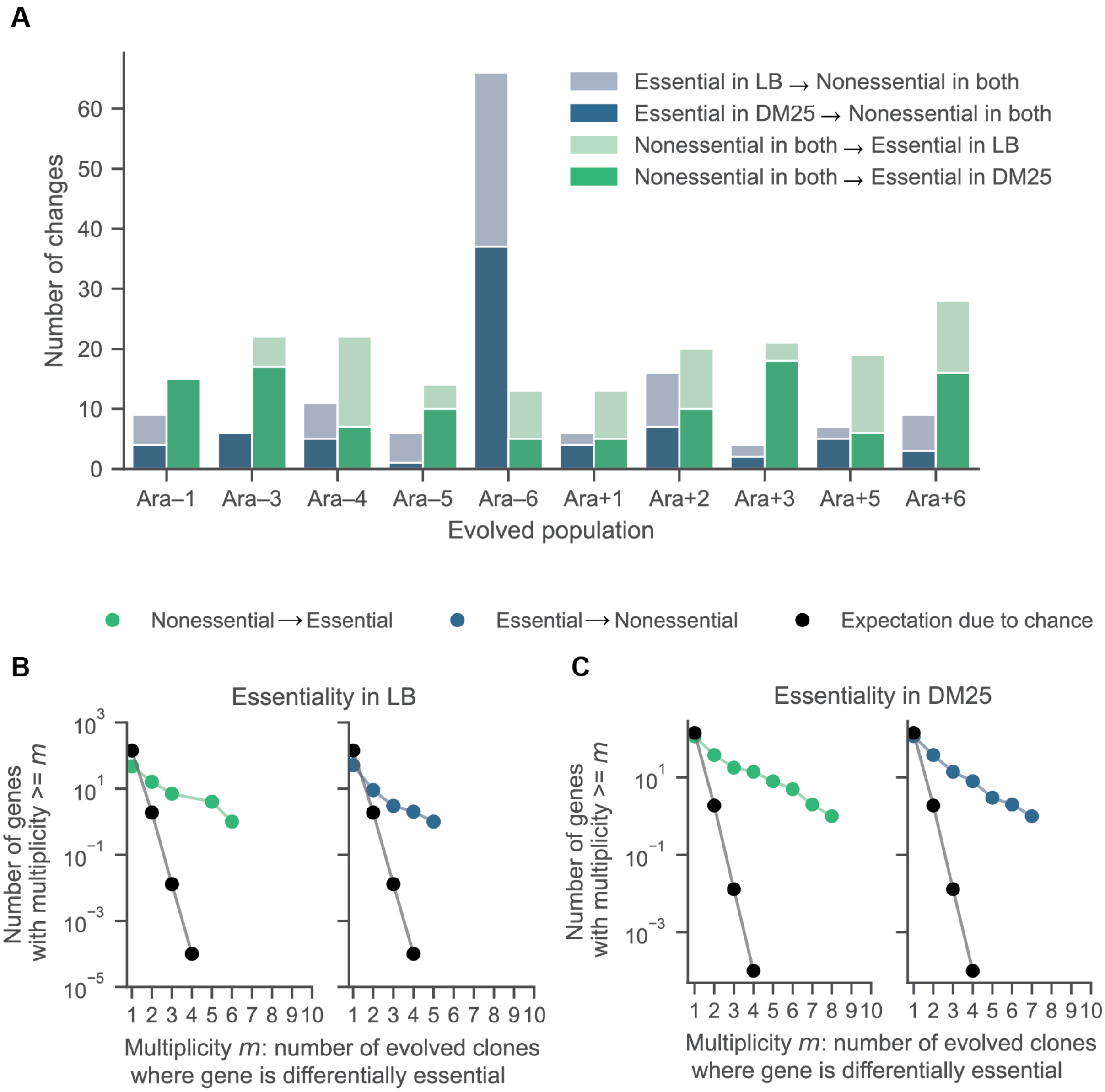

**Fig. S9: Parallelism in gene essentiality changes, partitioned by two different media. (A)** Partitioning changes in essentiality by growth medium. Parallel changes in gene essentiality in **(B)** LB and **(C)** DM25 media. We estimated the expected number of parallel changes from chance alone by shuffling the gene essentiality and fitness profiles 10,000 times and counting how often the same genes had altered essentially in at least *m* populations. The expectation is an average over 10,000 simulations, and therefore it can be < 1.

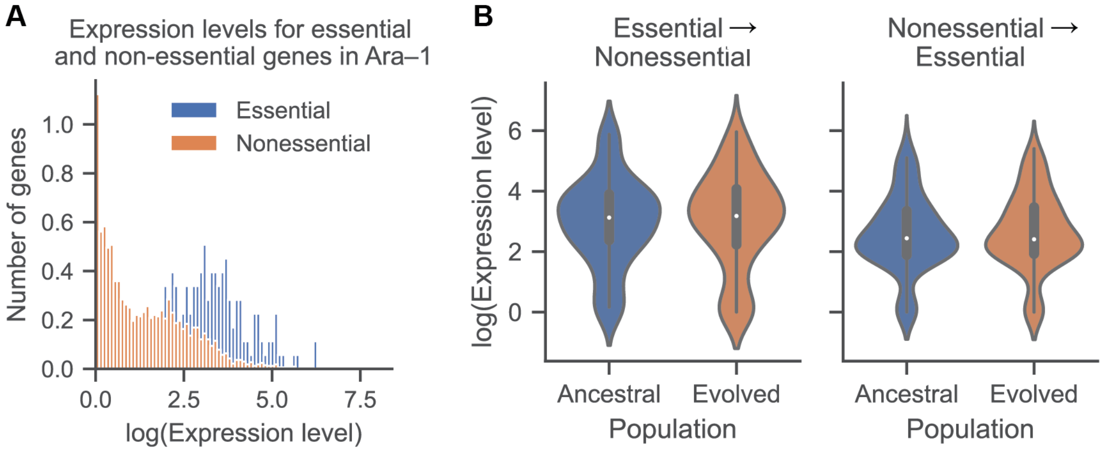

**Fig. S10: Changes in gene essentiality in DM25 are not related to differences in expression levels. (A)** Nonessential genes tend to have lower expression levels. The pattern shown for Ara–1 occurs in all LTEE clones (see Jupyter Notebook generate_figures_main.ipynb section on expression levels comparison). **(B)** Changes in gene essentiality do not reflect systematic differences in expression levels between the ancestor and evolved strains. Here, we show all genes with altered essentiality in any evolved strain. We compare the expression level of the gene in the ancestor and the evolved strain with altered essentiality for all these genes. The baseline expression levels are generally higher in the evolved strains; when comparing strains, we therefore normalized the expression levels of each gene by the total number of reads that mapped to the strain.

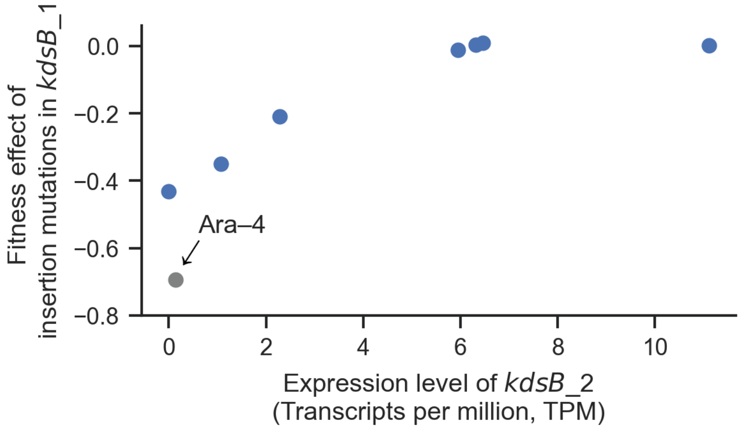

**Fig. S11: Fitness effect of disrupting *kdsB_1* gene is correlated with expression (transcripts per million, TPM) of *kdsB_2*.** A fitness effect could not be directly calculated for the outlier Ara–4 strain, owing to the mutants' rapid loss from the bulk competition assay. Pearson $r = 0.884$, $p = 0.0083$ excluding Ara–4; Pearson $r = 0.714$, $p = 0.0083$ with Ara–4 set to maximum deleterious effect of $-\ln(100)/\log_2(100) \cong -0.693$.

### List of Supplementary Tables:

**Table S1:** Counts from fitness assays based on direct competitions

**Table S2:** List of all genes differentially essential over evolution

**Table S3:** Homologous gene groups in the ancestral strain ANC

**Table S4:** Fitness effects of *wecA* insertion mutants

**Table S5:** Essentiality and expression levels of *kdsB* genes

### Detailed Experimental Protocols:

#### *Part 1: Constructing Transposon Libraries*

NOTE: This protocol is adapted from one provided by Thao Truong in the Thomas Bernhardt Lab in the Department of Microbiology at HMS. Many thanks to Thao for sharing this protocol and helping with initial troubleshooting.

Materials

Use DAP at final concentration of 300 uM in plates and liquid media

- Sigma 33240-1G (2,6-Diaminopimelic acid))
- Make 60 mM solution in 50 mL water, dissolve by vigorous stirring, store at 4C

Millipore MFilters

- 0.45 um catalog #HAWP02500

Regular sterile petri dishes (100 mm diameter)
Large sterile petri dishes (150 mm diameter)

Donor Strain

Freshly transform competent donor MFDλpir with plasmid pSC189 for maximum efficiency, and then select on LB agar plates supplemented with Kan 50 ug/mL). The plasmid has both Kan and Amp resistance genes, but selection on Kan was more effective.

Transformants were occasionally heterogeneous in size, and some were mucoid-like. We used colonies that were non-mucoid and sometimes small without problems.

Optimization

After running some simulations on the effect of library complexity (i.e., how many unique colonies one would expect after conjugation reactions), we found that complexity should have little effect on the precision of fitness estimates. That result occurs because, given a fixed number of reads, there is a tradeoff between having more reads for fewer TA sites, on the one hand, and fewer reads per TA site in each gene. In our experiments, we aimed to produce libraries with a complexity of at least 250,000. Such complexity is crucial for analysis of gene essentiality but is not strictly required for accurate fitness estimates.

To increase library complexity:

- Use freshly prepared MFDλpir/pSC189 (i.e., within 4-5 days of transformation)
- Scale up accordingly. For some genetic backgrounds, we increased the number of large plates on which we spread the libraries. Achieving high-complexity libraries was not a problem for most of the LTEE isolates, with the exception of the 50,000-generation clone REL11370 from population Ara+6.

Day 1

Streak relevant strains on agar plates

- MFDlpir/pSC189 on LB with DAP and Kan 50 (ug/mL)
- Recipient strains on LB

Day 2

Start 2-3 mL overnight cultures of each strain. A total of 5 mL of the donor strain is sufficient for two recipient strains. If more is required (e.g., more than two recipients), scale up the number of donor cultures. Grow the cultures either shaking or rolling at 37 C overnight, with ~16 hours of growth at most.

Day 3

Wash cells

1. For each strain, spin 1 mL of culture at 15,000 rpm for 1 min in Eppendorf tubes. Pellet 1 mL of donor cells for each conjugation.
2. Remove supernatant
3. Resuspend in 1 mL LB + DAP to wash cells
4. Spin again at 15,000 rpm for 2 min

Mate cells

1. Sterilize tweezers in 70% ethanol and then flame
2. For each mating (donor + recipient) transfer a filter to a plate of normal LB + DAP
3. Multiple filters can be placed on one plate as long as the filters are not touching
4. For each control (donor only, recipient only), transfer 1 filter to a plate of normal LB + DAP

Mate donor + recipient cells

1. Resuspend donor cells from one Eppendorf tube in ~100 μL of LB DAP
2. Transfer this volume to the Eppendorf tube containing the pelleted recipient, resuspend the cell mixture, and transfer them to the filter
3. Repeat this procedure for each recipient

For the controls, plate the donors or recipients alone directly on LB + Kan 50 agar plates.

Working quickly, transfer the mating plates (including controls) to a 37 C incubator or room, keeping the plates upright to avoid disturbing the filters. Incubate the plates upright (i.e., with filters facing up) for only 1 h.

Plate libraries

1. During incubation, prepare 15-mL conical tubes with media for plating steps, and 1.5-mL tubes for serial dilutions
2. Sterilize tweezers in 70% ethanol and flame
3. Transfer each filter from mating into a 15-mL conical tube containing 2 mL of LB + Kan 50 without DAP. This medium is the same as subsequently used for plating libraries, except the plating medium also has agar. The liquid volume can be increased if the multiple filters are used
4. Vortex the tube for 1 min
5. Serially dilute 100 μL from the tube across 4 successive 1.5-mL tubes, each containing 900 μL LB + Kan
6. Spread 100 μL from the second, third, and fourth dilutions onto regular size plates that contain LB + Kan. Given the plating volume, these effectively become $10^{-3}$, $10^{-4}$, and $10^{-5}$ dilutions when calculating the library complexity.

7. From the undiluted mating mixture (from step 2), spread 300 μL each onto 3 large LB+Kan plates.
8. Incubate all plates at 30 C for ~18-24 hours.

NOTE: the incubation duration is sufficient for colonies to form on the plates during mating, while avoiding colonies on the donor-only and recipient-only controls

Day 4

Calculate the library complexity (number of unique colonies in the transposon library) from the dilution series and the total volume plated on large LB+Kan plates (for 3 plates, the total volume is 900 μL.

A successful experiment typically has library complexity of ~$10^6$ . If complexity is lower:

1. Increase the number of LB+Kan plates, and therefore the volume plated
2. Increase the volume of recipient used in a mating reaction from 1 mL to 1.5-2.5 mL

Freeze library aliquots

1. Scrape cells from the large LB+Kan plates using a cell spreader and ~4 mL of LB per plate.
2. After scraping all plates, pool the resulting cells.
3. Add glycerol from a 50% solution such that the final concentration of glycerol for the transposon library was ~17%.
4. Vortex the libraries and transferred 0.5-mL aliquots into 1.5-mL Eppendorf tubes.
5. Freeze aliquots at –80 C.

Estimate CFUs and confirm exconjugants have transposon

1. Scrape ~50 μL from a frozen aliquot and serially diluted in LB.
2. Spread 100 μL of $10^{-5}$, $10^{-6}$, $10^{-7}$ dilutions on LB plates to determine viable counts. We expected ~$10^{9}$-$10^{10}$ CFU/mL, but some libraries yielded lower numbers because those LTEE isolates do not grow as well on LB.
3. The next day, count colonies and patch~100 colonies on to LB and LB + Kan plates to verify that all clones are KanR, as expected if they harbor the transposon. Patch a KanS colony as a control.
4. If all, or at least the vast majority (>90%), of tested colonies are *not* KanR, then redo the experiment.

Troubleshooting inefficient kanamycin selection:

- For some genetic backgrounds, the conjugation is highly efficient, and too many cells are plated, leading to inefficient kanamycin selection. One approach is to reduce how many cells are plated on Kan. A rule of thumb is to reduce the volume plated based on what fraction of patched colonies are KanS
- If that does not resolve the low %KanR colonies, plate the cells on LB+Kan100 agar plates after the conjugation step. This does not affect growth of the conjugants in any way (the KanR colonies grow normally up to a concentration of Kan500), but significantly improves the efficiency of selection.

***Part 2: Fitness Assays***

Media and growth conditions:

All fitness assays are performed in DM25 (Davis-Mingioli minimal medium). The following recipe makes one 1000-ml batch. Note also that *E. coli* cannot use citrate to support growth; it serves only as a chelating agent in this medium.

1. Potassium phosphate dibasic trihydrate (Sigma P5504)- 7 g
2. Potassium phosphate monobasic anhydrous (Sigma P5379)- 2 g
3. Ammonium sulfate (EMD AX1385) - 1 g
4. Sodium citrate (Sigma C7254) - 0.5 g
5. Water - 1000 ml

Immediately after autoclaving, add the following from sterile stock solutions (kept in 100-ml bottles covered in foil)

1. 10% Glucose - 250 µL (Stock solution: 10 g glucose (Sigma G8270)in 100 mL water autoclaved and covered in foil)
2. 10% Magnesium sulfate - 1000 µL (Stock solution: 10 g MgSO4 (Sigma M7506) in 100 mL water autoclaved and covered in foil)
3. 0.2% Thiamine/Vitamin B1) - 1000 µL (0.2 g thiamine hydrochloride (Sigma T4625) in 100 mL water filter sterilized; do NOT autoclave)

All fitness assays are performed in 15mm glass test tubes, and incubated at 37C and 220rpm

*NOTE*: While this is different from the traditionally used Erlenmeyer Flasks, from personal communication with Tanush Jagdish (Murray and Desai Labs at Harvard), any well-mixed glass container is a good substitute. We decided to use glass tubes for the convenience of being able to do many fitness assays in parallel.

Daily Transfers:

1. For each transposon library, estimate the density of cells (CFU/mL in the frozen library)
2. Extract DNA from at least 5x10$^{8}$ CFU/mL thawed transposon library. This serves as time-point 0.
3. Seed 5 glass test tubes with 10mL of DM25 with ~5x10$^{6}$ cells from the transposon library for one fitness assay
4. Incubate and 37C at 220 rpm for 24 hours.
5. Pool the five cultures corresponding to the same fitness assay into a 50mL falcon tube. Mix thoroughly by vortexing.
6. Transfer 0.5mL of the pooled cultures into 49.5mL of DM25 and mix well by vortexing
7. Pellet the remaining overnight culture by centrifuging at 4300 rpm for 30 mins, removing most of the supernatant, suspending in 1ml of DM25, and centrifuging at 16,000 rpm for 2 mins.
8. Store the cell pellets at -20C. Call this time-point 1.
9. Pipette 10mL of the diluted cultures into labelled glass test tubes and incubate again at 37C and 220 rpm for 24 hours

10. Repeat steps 3-8 until time-point 4.

NOTE: By having 5 cultures per fitness assay, we increase the population size by a factor of 5, reducing noise from the daily dilution of cells into fresh media. We still continue to grow the cultures in 10mL volumes to replicate the conditions of LTEE as closely as possible, and avoid any potential density dependent effects in fitness measurements.

***Part 3: UMI-TnSeq Sequencing Library Preparation Protocol***

Step 1: DNA Extraction

1. Use the Invitrogen PureLink gDNA extraction kit to extract gDNA from upto ~2x$10^9$ cells (usually 1mL overnight culture in LB has this many cells, ~50 mL of overnight culture for LTEE isolates)
2. Measure concentration of gDNA using Invitrogen Quant-It Kit
3. Normalize concentrations of different preps to ~20 ng/µL

Step 2: Nextera Tagmentation Reaction

1. (for 10 µL reactions) Mix 5 µL buffer, 2.5 µL TDE1 enzyme, 2.5 µL of gDNA (approximately 50ng) in a PCR tube. Vortex briefly and pulse centrifuge the reaction mixture.
2. Incubate at 55°C for 10 mins in a thermocycler, keep on ice right after (or in the fridge at 4C if not doing the PCRs instantly)

**NOTE**: this protocol uses Nextera TDE1 Tagmentase enzyme, but in principle, this can be adapted to the Illumina Library Prep Kit which uses a bead-linked transposome.

Step 3: PCR 1 - adding unique molecular identifiers

1. Make 10 µM working stocks of all primers (listed in Part 4).
2. Make a list of barcode combinations for each sample (ensure that the barcodes are color balanced). The barcode combinations used in this study are shown in Part 5.
3. Aliquot 3 µL each of forward (TnSeq_1F) and reverse primer (TnSeq_1R_xx; choose the PCR 1 reverse primer according to the barcode combinations you've determined in step 2) in PCR tubes, add all 10 µL of the Nextera tagmentation reaction (serves as template).
4. Add 15 µL of the Q5 master mix and mix with pipette. Vortex briefly and pulse centrifuge the reaction mixture
5. Run the following PCR program:

| | | |
|---|---|---|
| 1 | 95°C | 3:00 |
| 2 | 98°C | 0:30 |
| 3 | 68°C | 0:30 |
| 4 | 72°C | 1:00 |
| 5 | Go to 2 | 2X |
| 6 | 72 °C | 3:00 |
| 7 | 12°C | hold |

Step 4: SPRI Bead Cleanup

1. Perform magnetic bead cleanup using ~1.2X serapure beads (home-brewed, the exact ratio will vary for AmpPure XP beads)
2. Elute in 15 µL dH$_2$0

Step 5: PCR 2 - amplification and addition of Illumina barcodes and adapter sequences

1. Make 10 µM working stocks of all primers.
2. Aliquot 5 µL each of forward (TnSeq_2F_yy; choose the PCR 2 forward primer according to the barcode combinations you've determined in step 2) and reverse primer (TnSeq_2R) in PCR tubes, add all 15 µL of the cleaned up PCR1 product
3. Add 25 µL Q5 master mix and pipette up and down to mix. Vortex briefly and pulse centrifuge the reaction mixture
4. Run the following PCR program:

| | | |
|---|---|---|
| 1 | 95°C | 3:00 |
| 2 | 98°C | 0:30 |
| 3 | 68°C | 0:30 |
| 4 | 72°C | 1:00 |
| 5 | Go to 2 | 18X |
| 6 | 72 °C | 3:00 |
| 7 | 12°C | hold |

Step 6: SPRI Bead Cleanup

1. Perform magnetic bead cleanup using ~1.2X serapure beads (home-brewed, the exact ratio needs to varied if you're using AmpPure XP beads)
2. Elute in 25 µL dH$_2$0

Step 7: Quantifying Sequencing Libraries

1. Use QuantIT Kit (or an equivalent) to estimate concentration of the libraries after PCR2.
2. Run a few of the libraries on a 2% Agarose Gel, with a 50bp ladder, and get an approximate sense of the average fragment size
3. Dilute and all libraries to approximately 4nM and pool the libraries together.
4. Perform a qPCR (KAPA Quantit Kit or an equivalent) on the pooled libraries. Make sure to run two dilutions (1:10,000, 1:20,000)
5. Calculate the concentration of the pooled library using the calibration curve.
6. Run the pooled library on an Agilent BioA high sensitivity DNA chip to get a more accurate sense of fragment size distribution.

7. Ensure that there are no primer dimers. With the current protocol, primer dimers are expected at ~125bp in case of unsuccessful bead clean-up. Re-calibrate your beads and ensure they're size selective if you observe primer dimers.
8. Dilute libraries to 4nM (or the appropriate concentration for the sequencing platform)

**Bead Clean-up Protocol:**

The bead clean-up step is most effectively done in PCR strips with a compatible magnetic rack, using a multichannel pipette.

1. Suspend sample in 1.2x the volume of beads, and mix. Incubate at room temperature for 10 mins
2. Place on magnetic rack, allow beads to pellet and pipette off supernatant
3. Add 100 µL of freshly prepared 70% ethanol
4. Move the PCR strip to adjacent location (this will move the beads through the ethanol, washing them)
5. Remove the supernatant.
6. Repeat steps 3-5
7. Leave tube open on the bench and let the ethanol evaporate
8. Remove the tube from the magnetic rack and resuspend pellet in an appropriate volume of eluant (e.g. 20 µL molecular grade water)
9. Incubate for 10 mins at room temperature
10. Pellet the beads on the magnetic stand until the eluate is clear. Transfer the supernatant to fresh PCR strips or eppendorfs.

### Part 4: Primers for UMI-TnSeq Library Preparation

| | |
|---|---|
| TnSeq_1R_01 | CAAGCAGAAGACGGCATACGAGAT**GGCGAATG**GTCTCGTGGGCTCGGAGAT |
| TnSeq_1R_02 | CAAGCAGAAGACGGCATACGAGAT**CGATAGAG**GTCTCGTGGGCTCGGAGAT |
| TnSeq_1R_03 | CAAGCAGAAGACGGCATACGAGAT**TTGCGTCA**GTCTCGTGGGCTCGGAGAT |
| TnSeq_1R_04 | CAAGCAGAAGACGGCATACGAGAT**ATGATTAA**GTCTCGTGGGCTCGGAGAT |
| TnSeq_1R_05 | CAAGCAGAAGACGGCATACGAGAT**TCTTCATC**GTCTCGTGGGCTCGGAGAT |
| TnSeq_1R_06 | CAAGCAGAAGACGGCATACGAGAT**TCAGTACG**GTCTCGTGGGCTCGGAGAT |
| TnSeq_1R_07 | CAAGCAGAAGACGGCATACGAGAT**CATACCTC**GTCTCGTGGGCTCGGAGAT |
| TnSeq_1R_08 | CAAGCAGAAGACGGCATACGAGAT**GGCCAGAA**GTCTCGTGGGCTCGGAGAT |
| TnSeq_1R_09 | CAAGCAGAAGACGGCATACGAGAT**GGTCAAGT**GTCTCGTGGGCTCGGAGAT |
| TnSeq_1R_10 | CAAGCAGAAGACGGCATACGAGAT**AACTCTCT**GTCTCGTGGGCTCGGAGAT |
| TnSeq_1R_11 | CAAGCAGAAGACGGCATACGAGAT**CTAGGTTA**GTCTCGTGGGCTCGGAGAT |
| TnSeq_1R_12 | CAAGCAGAAGACGGCATACGAGAT**TCCGGCGG**GTCTCGTGGGCTCGGAGAT |
| TnSeq_1R_13 | CAAGCAGAAGACGGCATACGAGAT**ACGATCGC**GTCTCGTGGGCTCGGAGAT |
| TnSeq_1R_14 | CAAGCAGAAGACGGCATACGAGAT**GTTATGCT**GTCTCGTGGGCTCGGAGAT |
| TnSeq_1R_15 | CAAGCAGAAGACGGCATACGAGAT**CAGTCGGT**GTCTCGTGGGCTCGGAGAT |
| TnSeq_1R_16 | CAAGCAGAAGACGGCATACGAGAT**AACCGCAA**GTCTCGTGGGCTCGGAGAT |
| TnSeq_1F | CTCTTTCCCTACACGACGCTCTTCCGATCT**AANNNNNTTGGGGACTTATCAGCCAACCT** |
| TnSeq_2F_01 | AATGATACGGCGACCACCGAGATCTACAC**GGACGTAG**ACACTCTTTCCCTACACGACGCT |
| TnSeq_2F_02 | AATGATACGGCGACCACCGAGATCTACAC**GATGCCAG**ACACTCT |

| | |
|---|---|
| | TTCCCTACACGACGCT |
| TnSeq_2F_03 | AATGATACGGCGACCACCGAGATCTACAC**TCGGATGC**ACACTCT TTCCCTACACGACGCT |
| TnSeq_2F_04 | AATGATACGGCGACCACCGAGATCTACAC**ATAACTTC**ACACTCT TTCCCTACACGACGCT |
| TnSeq_2F_05 | AATGATACGGCGACCACCGAGATCTACAC**TTCTGGCT**ACACTCT TTCCCTACACGACGCT |
| TnSeq_2F_06 | AATGATACGGCGACCACCGAGATCTACAC**CCGGTAAC**ACACTCT TTCCCTACACGACGCT |
| TnSeq_2F_07 | AATGATACGGCGACCACCGAGATCTACAC**CAATGCCT**ACACTCT TTCCCTACACGACGCT |
| TnSeq_2F_08 | AATGATACGGCGACCACCGAGATCTACAC**AGATAAGA**ACACTCT TTCCCTACACGACGCT |
| TnSeq_2F_09 | AATGATACGGCGACCACCGAGATCTACAC**CCTAATCT**ACACTCT TTCCCTACACGACGCT |
| TnSeq_2F_10 | AATGATACGGCGACCACCGAGATCTACAC**CATCTCCG**ACACTCT TTCCCTACACGACGCT |
| TnSeq_2F_11 | AATGATACGGCGACCACCGAGATCTACAC**ACGCAGTC**ACACTCT TTCCCTACACGACGCT |
| TnSeq_2F_12 | AATGATACGGCGACCACCGAGATCTACAC**TTCCTCTA**ACACTCT TTCCCTACACGACGCT |
| TnSeq_2F_13 | AATGATACGGCGACCACCGAGATCTACAC**GGTTCGTA**ACACTCT TTCCCTACACGACGCT |
| TnSeq_2F_14 | AATGATACGGCGACCACCGAGATCTACAC**GTAACGAG**ACACTCT TTCCCTACACGACGCT |
| TnSeq_2F_15 | AATGATACGGCGACCACCGAGATCTACAC**AAGAGAGT**ACACTCT TTCCCTACACGACGCT |
| TnSeq_2F_16 | AATGATACGGCGACCACCGAGATCTACAC**TGCGTAGC**ACACTCT TTCCCTACACGACGCT |
| TnSeq_2R | CAAGCAGAAGACGGCATACGA |

### Part 5: Primer Combinations for Samples:

| | primer combination | | | | | | | | |
|---|---|---|---|---|---|---|---|---|---|
| format (x,y) | (1R_x, 2F_y) | | | Fitness assay time point | | | | | |
| genetic background | t0 | t1 | t1 | t2 | t2 | t3 | t3 | t4 | t4 |
| REL606 | (1,1) | (2,1) | (3,1) | (4,1) | (5,1) | (6,1) | (7,1) | (8,1) | (9,1) |
| REL607 | (1,2) | (2,2) | (3,2) | (4,2) | (5,2) | (6,2) | (7,2) | (8,2) | (9,2) |
| REL11330 (Ara–1) | (1,3) | (2,3) | (3,3) | (4,3) | (5,3) | (6,3) | (7,3) | (8,3) | (9,3) |
| REL11333 (Ara–2) | (1,4) | (2,4) | (3,4) | (4,4) | (5,4) | (6,4) | (7,4) | (8,4) | (9,4) |
| REL11364 (Ara–3) | (1,5) | (2,5) | (3,5) | (4,5) | (5,5) | (6,5) | (7,5) | (8,5) | (9,5) |
| REL11336 (Ara–4) | (1,6) | (2,6) | (3,6) | (4,6) | (5,6) | (6,6) | (7,6) | (8,6) | (9,6) |
| REL11339 (Ara–5) | (1,7) | (2,7) | (3,7) | (4,7) | (5,7) | (6,7) | (7,7) | (8,7) | (9,7) |
| REL11389 (Ara–6) | (1,8) | (2,8) | (3,8) | (4,8) | (5,8) | (6,8) | (7,8) | (8,8) | (9,8) |
| REL11392 (Ara+1) | (1,9) | (2,9) | (3,9) | (4,9) | (5,9) | (6,9) | (7,9) | (8,9) | (9,9) |
| REL11342 (Ara+2) | (1,10) | (2,10) | (3,10) | (4,10) | (5,10) | (6,10) | (7,10) | (8,10) | (9,10) |
| REL11345 (Ara+3) | (1,11) | (2,11) | (3,11) | (4,11) | (5,11) | (6,11) | (7,11) | (8,11) | (9,11) |
| REL11348 (Ara+4) | (1,12) | (2,12) | (3,12) | (4,12) | (5,12) | (6,12) | (7,12) | (8,12) | (9,12) |
| REL11367 (Ara+5) | (1,13) | (2,13) | (3,13) | (4,13) | (5,13) | (6,13) | (7,13) | (8,13) | (9,13) |
| REL11370 (Ara+6) | (1,14) | (2,14) | (3,14) | (4,14) | (5,14) | (6,14) | (7,14) | (8,14) | (9,14) |

**Part 6: Primers for Recombineering:**

| | |
|---|---|
| **recomb_insert_reverse** | **CTTTCTACGTGTTCCGCTTC** |
| nadR_recomb_confirm_f | CTAACGTACCGAAAAAAGCGC |
| nadR_recomb_KanR_f | GCGTGTTCGACGACTTATAATGAGGAATACGGAGGGAGA T**GTGTAGGCTGGAGCTGCTT** |
| nadR_recomb_KanR_r | AACTTATTTTATACCTCGCCTTTGAGCCGTTTCATCGCGG**C ATATGAATATCCTCCTTA** |
| nadR_recomb_confirm_r | CCCGTGCAGGTTTTTTCAAAC |
| rbsD_recomb_confirm_f | CGGATAGACATTTAACGCTGC |
| rbsD_recomb_KanR_f | ATCGATGCCTTTAAGCTGAAGTAATGCTTCCATGACGGCC **GTGTAGGCTGGAGCTGCTT** |
| rbsD_recomb_KanR_r | AACTGTGGGTCAGCGAAACGTTTCGCTGATGGAGAAAAA A**CATATGAATATCCTCCTTA** |
| rbsD_recomb_confirm_r | TTTCATGGTTAATCACCATGTAAAACG |
| lacA_recomb_confirm_f | GTGGGTCAAAGAGGCATGATG |
| lacA_recomb_KanR_f | CAGCGTATCAGGCAATTTTTATAATTTAAACTGACGATTC **GTGTAGGCTGGAGCTGCTT** |
| lacA_recomb_KanR_r | AACATATCAGGACGGAATGATCGCAATGAACATGCCAAT G**CATATGAATATCCTCCTTA** |
| lacA_recomb_confirm_r | CTTAATTTCCGTGTTCACGCTTAG |
| dctA_recomb_confirm_f | CATGAAAATGTCACGGAAGAAGTGA |
| dctA_recomb_KanR_f | CCCGCACTCGGGGAAGGGAGTGCGGGCATAAGTGATGAG A**GTGTAGGCTGGAGCTGCTT** |
| dctA_recomb_KanR_r | ATAACCTTACAAGATCTGTGGTTTTACTAAAGGACACCCT **CATATGAATATCCTCCTTA** |
| dctA_recomb_confirm_r | GCTCAAACTTTTTAACCTTTTTGTTTCAAT |
| gdhA_recomb_confirm_f | TTGCTTTCCTGGGTCATTTTTTTCTT |
| gdhA_recomb_KanR_f | CATAAGCACAATCGTATTAATATATAAGGGTTTTATATCT **GTGTAGGCTGGAGCTGCTT** |
| gdhA_recomb_KanR_r | AGCGTAGCGCCATCAGGCATTTACAACTTAAATCACACCC **CATATGAATATCCTCCTTA** |
| gdhA_recomb_confirm_r | AATACTCATAAACGCCTGAAATTTTGC |
| fecE_recomb_confirm_f | CGGATGGACGATGAGTATTGGTAA |
| fecE_recomb_KanR_f | TTTCATTCAGTCGTGGTTTGGTTCTTACGGCCTGTGCAAT**G TGTAGGCTGGAGCTGCTT** |
| fecE_recomb_KanR_r | TGCGCCGTGGTTTGTCTGGTTGCTTGTGAGAATGCGATAA |

| | |
|---|---|
| | **CATATGAATATCCTCCTTA** |
| fecE_recomb_confirm_r | CGATCTGCTGGCGAGAATTAT |
| ydiJ_recomb_confirm_f | TACTGGCATTGTCGCTTCAAGG |
| ydiJ_recomb_KanR_f | ATAGCATTCAGTGCTTCCAGGGTGATTTTCCGTTTCCATA**GTGTAGGCTGGAGCTGCTT** |
| ydiJ_recomb_KanR_r | TATCGACCTACATCACAGACCGCAGGAAAGGGTCAATATA **CATATGAATATCCTCCTTA** |
| ydiJ_recomb_confirm_r | GCCTTCCTGCAACACGAAAT |
| sspA_recomb_confirm_f | GCATATTCCATAGGAACCTGCAC |
| sspA_recomb_KanR_f | TAGGGACGACGTGGTGTTAGCTGTGACAAATCCATACAGA **GTGTAGGCTGGAGCTGCTT** |
| sspA_recomb_KanR_r | CTGGTAGCAGTAAAAATTCTGACTATACCTGGAGGTTTTC **CATATGAATATCCTCCTTA** |
| sspA_recomb_confirm_r | GGGTATTGCTCATTTTTTGTTTGATT |
| proC_recomb_confirm_f | CCAAATTGTCATAAAGTCATCCTTTGTT |
| proC_recomb_KanR_f | AACAATGAATTTCACGGCAGGAGTGAGGCAATGGAAAGA**GTGTAGGCTGGAGCTGCTT** |
| proC_recomb_KanR_r | AACCGCACCGAAGTGGCGGCCTGACGTCCGGCGAAAGTCA**CATATGAATATCCTCCTTA** |
| proC_recomb_confirm_r | GCTGAACCCACAAATGAGTCAC |
| ppc_recomb_confirm_f | CCTTAAGGATATCTGAAGGTATATTCAGAATTTG |
| ppc_recomb_KanR_f | ACCCTCGCGCAAAAGCACGAGGGTTTGCAGAAGAGGAAGA**GTGTAGGCTGGAGCTGCTT** |
| ppc_recomb_KanR_r | ACAGGGCTATCAAACGATAAGATGGGGTGTCTGGGGTAAT**CATATGAATATCCTCCTTA** |
| ppc_recomb_confirm_r | CACCGCTTTTACGTGGCTTTAT |
